# The cGAS-STING Pathway Is Essential in Acute Ischemia-Induced Neutropoiesis and Neutrophil Priming in the Bone Marrow

**DOI:** 10.1101/2024.07.18.604120

**Authors:** Jiankun Zhu, Xinjia Ruan, MariaSanta C Mangione, Pablo Parra, Xiaoping Su, Xiang Luo, Dian J Cao

## Abstract

Acute myocardial ischemia triggers a rapid mobilization of neutrophils from the bone marrow to peripheral blood, facilitating their infiltration into the infarcted myocardium. These cells are critical for inducing inflammation and contributing to myocardial repair. While neutrophils in infarcted tissue are better characterized, our understanding of whether and how ischemia regulates neutrophil production, differentiation, and functionality in the bone marrow remains limited. This study investigates these processes and the influence of the cGAS-STING pathway in the context of myocardial infarction. The cGAS-STING pathway detects aberrant DNA within cells, activates STING, and initiates downstream signaling cascades involving NFKB and IRF3. We analyzed neutrophils from bone marrow, peripheral blood, and infarct tissues using MI models generated from wild-type, *Cgas*^-/-^, and *Sting*^-/-^ mice. These models are essential for studying neutropoiesis (neutrophil production and differentiation), as it involves multiple cell types. RNA sequencing analysis revealed that ischemia not only increased neutrophil production but also promoted cytokine signaling, phagocytosis, chemotaxis, and degranulation in the bone marrow before their release into the peripheral blood. Inhibition of the cGAS-STING pathway decreased neutrophil production after MI and down-regulated the same pathways activated by ischemia. Neutrophils lacking cGAS or STING were less mature, exhibited reduced activation, and decreased degranulation. Deletion of cGAS and STING decreased the expression of a large group of IFN-stimulated genes and IFIT1+ neutrophils from peripheral blood and the infarct tissue, suggesting that cGAS-STING plays an essential role in neutrophils with the IFN-stimulated gene signature. Importantly, transcriptomic analysis of *Cgas*^-/-^ and *Sting*^-/-^ neutrophils from bone marrow and MI tissues showed downregulation of similar pathways, indicating that the functionality developed in the bone marrow was maintained despite infarct-induced stimulation. These findings highlight the importance of neutropoiesis in dictating neutrophil function in target tissues, underscoring the critical role of the cGAS-STING pathway in neutrophil-mediated myocardial repair post-ischemia.

## Introduction

In acute myocardial ischemia, neutrophils are the first innate immune cells recruited into the damaged myocardium. They peak in abundance three days after myocardial infarction (MI) and are a dominant cell type in the ischemic tissue^1–4^. Neutrophils facilitate myocardial repair by dismantling necrotic tissue, removing tissue debris by phagocytosis, releasing matrix metalloproteases (MMPs) to remodel the provisional matrix, triggering inflammation, and attracting monocytes to the injury site^5^. At the same time, an overzealous response from these cells amplifies myocardial injury.

After MI and in other settings of systemic inflammation, hematopoietic stem and progenitor cells (HSPC) proliferate, commit to the granulocyte lineage, and then differentiate into mature neutrophils at a much faster rate^6, 7^. This process is known as "emergent neutropoiesis". During neutropoiesis, neutrophils pre-package tools they ultimately use to eliminate pathogens and trigger inflammation. This preparation starts when a progenitor cell commits to the neutrophil lineage and continues throughout the differentiation and maturation process in the bone marrow (BM). For example, primary, secondary, tertiary, and secretory granules containing enzymes and bactericidal molecules are sequentially generated during this process^8,9^. The expression of genes involved in the cell cycle and transcription/translation peaks early in differentiation and then decreases. At the same time, the expression of genes related to chemotaxis and response to microbial stimuli increases as neutrophils mature^10, 11^. At any stage of the differentiation continuum, neutrophils in the BM responded to infectious stimuli by enhancing cell migration, cytokine production, and granule formation^11^. These data suggest that inflammatory and infectious stimuli not only increase neutrophil numbers through neutropoiesis but also augment neutrophil functionality to prepare them for action at injury sites. Furthermore, single-cell omics have revealed that neutrophils are a diverse group of cells at baseline^11^, in different tissue environments^12–14^, and when responding to various infectious^11^ and inflammatory stimuli^12, 15, 16^. It remains unclear whether neutropoiesis contributes to this diversity. Thus, understanding neutropoiesis is crucial for comprehending neutrophil biology in specific pathological conditions. In acute myocardial ischemia, it remains unknown if and how this condition primes neutrophils during neutropoiesis in the BM, and whether such priming affects the phenotypes of neutrophils in the infarcted tissue.

At the transcriptomic level, one identified subtype of neutrophils exhibits an interferon-stimulated gene (ISG) expression signature. This subtype was found in peripheral blood and spleen at homeostasis and expanded in response to inflammatory stimuli^11^. ISG-related neutrophils were also identified in the infarcting myocardium^17, 18^. A recent study demonstrated that neutrophils had higher or similar ISG scores (the summation of 10 ISG expressions) compared to monocytes collected from the same compartments at the same time after MI, and the transcription factor IRF3 was key to the interferon response in these myeloid cells^19^. IRF3 activation occurs downstream of cGAS-STING-mediated signaling^20, 21^, which is also known to maintain a low or "tonic" level of IFN at homeostasis^22^. cGAS detects cytosolic double-stranded DNA (dsDNA) originating from pathogens or self-DNA. This dsDNA activates cGAS to synthesize cGAMP, a second messenger that activates STING. STING, in turn, activates transcription by IRF3 and NFKB, leading to the production of type I interferons (IFN-I), IFN-stimulated genes (ISGs), and other mediators of inflammation^20, 21^. The transcriptomic signature of ISG neutrophils includes hallmark genes of cGAS-STING activation, such as *Ifit1*, *Ifit3*, and *Irf7*^23, 24^. Thus, cGAS-STING may play a role in the generation and functions of the ISG-related neutrophils in acute myocardial ischemia. This role, however, has yet to be defined. Additionally, we and others have reported that cGAS-STING signaling is detrimental in post-myocardial infarction repair and remodeling by promoting inflammatory macrophages^23, 24^. Whether and how this pathway governs neutrophils in acute ischemia remains unknown. Few studies have investigated the cGAS-STING pathway in neutrophil differentiation and effector function. In fact, it remains to be determined whether neutrophils express cGAS.

Here, we investigated whether ischemia enhances neutrophil functionality and neutropoiesis in the bone marrow, and we examined neutrophil biology in cGAS- and STING-deficient mice at baseline and after MI. We discovered that acute ischemia induced broad-spectrum changes in BM neutrophils related to innate and adaptive immunity before releasing them into the peripheral blood. We found that the loss of cGAS and STING altered the transcriptomes of BM neutrophils, leading to the down-regulation of genes associated with cell migration, chemotaxis, cytokine response, and leukocyte activation. Additionally, we determined that cGAS-STING signaling was responsible for expressing a large group of ISGs in neutrophils after MI. Furthermore, the down-regulated transcriptomes observed in the BM were maintained in neutrophils in the infarct tissue, suggesting the functional relevance of neutrophil priming before infiltrating the ischemic injury site.

## Result

### Expression of cGAS-STING pathway components in neutrophils

Neutrophils were previously reported to express low levels of STING but not cGAS^25^. With the availability of cGAS-specific antibodies with improved sensitivity, we reassessed cGAS expression in neutrophils and also evaluated STING and IRF3 protein expression, the key nodes downstream of cGAS activation. We used three methods to purify neutrophils from WT, *Cgas*^-/-^, *Stin*g^-/-^, and *Irf3*^-/-^ mice. First, we isolated neutrophils by gradient centrifugation of bone marrow cells. The non-neutrophil phase was enriched for monocytes (labeled "Monocyte+") and used as a positive control for protein expression. BM neutrophils purified using this method expressed cGAS, STING, and IRF3 (**Figure 1A**). Negative controls using monocytes and neutrophils from *Cgas*^-/-^, *Stin*g^-/-^, and *Irf3*^-/-^ mice lacked the corresponding protein bands (**Figure 1A**). We performed flow cytometry after labeling the cells from the neutrophil phase with myeloid markers (CD45 and CD11b) and the neutrophil marker Ly6G to assess purity. **Figure 1B** demonstrates the highly purified neutrophil population by this method. Modified Giemsa staining also confirmed the high purity by identifying neutrophils based on their unique nuclear morphology (**Supplemental Figure**). Neutrophils exhibited lower cGAS and STING expression than monocytes but higher IRF3 levels (**Figure 1C**). Subcellular localization studies revealed that cGAS was primarily detected in the nuclear compartment of neutrophils at baseline (**Figure 1D**), similar to the cellular location of cGAS in macrophages^26^. Next, we purified BM neutrophils using fluorescence-activated cell sorting (FACS) or anti-Ly6G positive selection. FACS-purified BM neutrophils were gated as the CD45^+^CD11b^+^Ly6G^+^ population and labeled as such. The positively selected neutrophils were designated as Ly6G^+^ (**Figure 1E**). cGAS was detectable in BM neutrophils isolated by both methods (**Figure 1E**). STING was detected in Ly6G positively selected neutrophils (**Figure 1F**). However, immunoblots of lysates from FACS-purified BM neutrophils revealed a strong non-specific band at the same molecular weight as STING (data not shown). cGAS was also detected in cardiac neutrophils isolated from ischemic tissue three days after the LAD ligation procedure for MI (**Figure 1G**). In summary, our data demonstrate that neutrophils express cGAS and STING at baseline, though at lower levels than monocytes, especially for STING.

**Figure 1.**
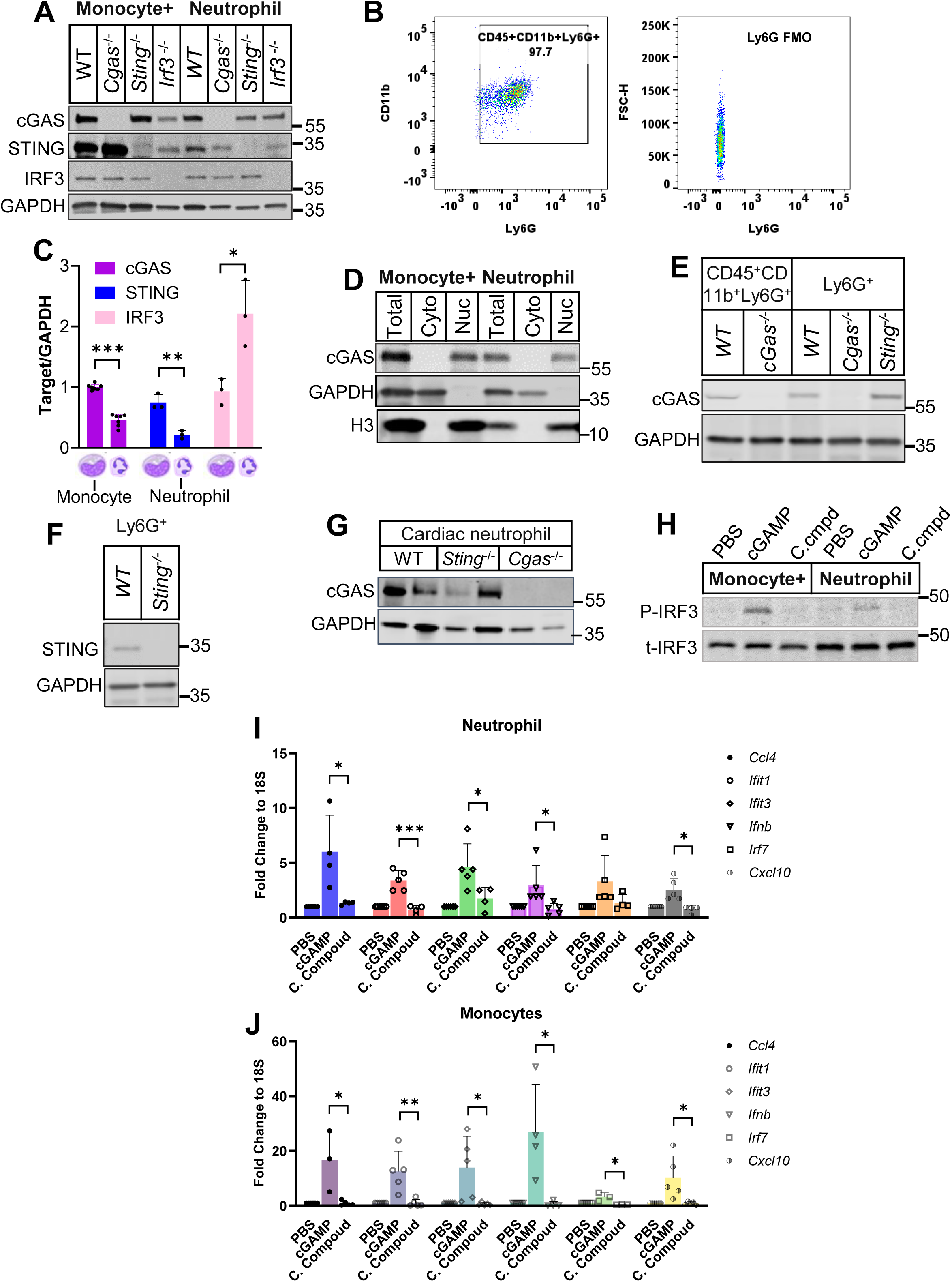
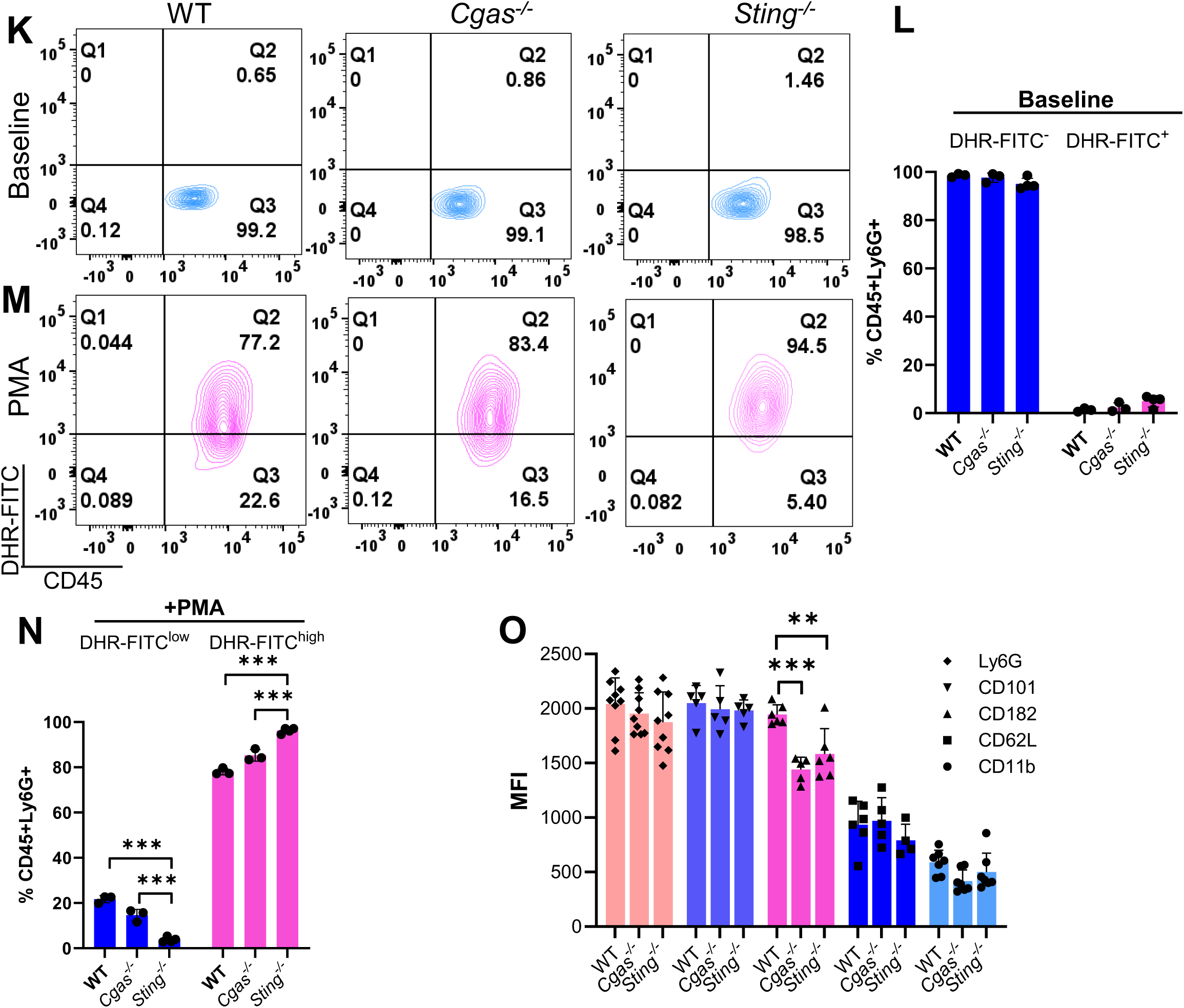
Neutrophils express components of the cGAS-STING pathway and baseline characterization of *Cgas*- and *Sting*-deficient neutrophils. Neutrophils were purified using three different methods. **A-D.** Bone marrow (BM) cells from mice with the indicated genotypes were separated by density gradient centrifugation into non-neutrophil and neutrophil phases. The non-neutrophil layer was monocyte-predominant and therefore labeled as “Monocyte+”. Cell lysates were probed using antibodies for the indicated protein (**A**). Neutrophil purity was assessed by flow cytometry after labeling the cells with CD11b, CD45, and Ly6G (**B**). Relative abundance of the indicated proteins as determined by immunoblotting as described in **A** (**C**). WT monocytes and neutrophils were fractionated to determine cGAS sub-cellular localization (**D**). **E**. Cell lysates from BM neutrophils of the indicated genotype purified by FACS using gate CD45+CD11b+Ly6G+ or Ly6G-beads positive selection (Ly6G+) were probed for cGAS. **F**. Cell lysates from BM neutrophils of the indicated genotype purified by Ly6G-beads positive selection (Ly6G+) were probed for STING. **G**. Neutrophils isolated from the heart 3 days after MI surgery were probed for the indicated protein. **H-J**. Neutrophils and monocytes from BM were stimulated with 12 μM cGAMP or its control compound (C.cmpd) for 90 min. Cells were used for immunoblot (**H**) or RT-qPCR (**I-J**). **K-L**. Peripheral blood

### Functional characterization of the cGAS-STING pathway in neutrophils

Activated cGAS-STING phosphorylates and activates IRF3. This molecular cascade eventually results in interferon-stimulated gene (ISG) expression. To determine if the cGAS-STING pathway is functional in neutrophils, we treated BM neutrophils isolated by gradient centrifugation with either cGAMP, a control compound for cGAMP (C. Compound or C.cmpd), or vehicle (PBS) for 90 min. The non-neutrophil layer, which was enriched in monocytes (Supplement Figure), served as the positive control. BM neutrophils treated with cGAMP had increased phosphorylated IRF3 (**Figure 1H).** Additionally, cGAMP treatment induced expression of *Ifnb* and ISGs, including *Ccl4, Ifit1*, *Ifit3, Irf7, and Cxcl10,* in BM neutrophils (**Figure 1I**) although at a reduced magnitude compared to monocytes (**Figure 1J)**, Thus, BM neutrophils are cGAMP-responsive, suggesting an intact cGAS-STING-IRF3 signaling cascade.

Next, we analyzed neutrophils from peripheral blood (PB) at baseline. We compared ROS production in WT, *Cgas^-/-^, and Sting^-/-^* PB neutrophils at homeostasis and after treatment with phorbol 12-myristate 13-acetate (PMA) by measuring the conversion of the cell-permeable dehydrorhodamine 123 (DHR123) to green-fluorescent rhodamine 123 (DHR-FITC^+^). At baseline, the percentage of DHR-FITC^+^ neutrophils was low and comparable among genotypes (**Figure 1K, 1L**), indicating the intrinsic ROS generated by neutrophils is minimal without stimulation. With PMA, the percentages of DHR-FITC^+^ neutrophils were similar between WT and *Cgas*-deficient neutrophils (**Figure 1M, 1N**), suggesting a similar capacity in generating ROS with stimulation. *Sting*^-/-^ neutrophils had a higher percentage of DHR-FITC^+^-neutrophils than WT and *Cgas*^-/-^ neutrophils (**Figure 1M, 1N)**.

To further determine if WT, *Cgas^-/-^*, and *Sting^-/-^* neutrophils differ at homeostasis, we compared the expression of cell surface markers of maturation (Ly6G, CD101), activation (CD11b), adhesion/migration (CD62L), chemotaxis (CD182), and degranulation (CD63) in PB neutrophils (CD45^+^CD11b^+^Ly6G^+^). We used median fluorescent intensity (MFI) to indicate the levels of these markers and found that Ly6G, CD101, CD11b, and CD62L were similar among WT, *Cgas^-/-^, and Sting^-/-^* neutrophils at baseline. However, CD182 was significantly decreased by 20-25% in neutrophils from *Cgas^-/-^* and *Sting^-/-^* mice (**Figure 1O**). CD182 (CXCR2) is a G protein-coupled receptor crucial for neutrophil chemotaxis and trans-epithelial migration. This reduction in CD182 at steady state did not affect the neutrophil infiltration in the MI tissue, which will be discussed later, suggesting a limited impact on neutrophil function. In summary, while cGAS-STING signaling was intact in BM neutrophils, loss of *Cgas* or *Sting* had minimal effects on PB neutrophil function at baseline; ROS production was preserved (slightly increased in *Sting^-/-^*), and markers of maturation, activation, adhesion, migration, and degranulation were largely similar. CD63 was not detectable in these neutrophils (not shown), which is expected in WT cells, suggesting the degranulation process was not perturbed in baseline conditions by *Cgas* and *Sting* deletion.

### cGAS-STING promotes neutrophil production and maturation in the bone marrow after myocardial infarction

We next investigated whether cGAS-STING pathway impacts neutrophils in the setting of myocardial infarction (MI). First, we measured the neutrophilic response three days after MI using the neutrophil percentage of CD45^+^ white blood cells in the peripheral blood (PB). MI induced a 3-fold increase in circulating neutrophils, which was comparable among *WT, Cgas*^-/-^, and *Stin*g^-/-^ mice (**Figure 2A, 2B**).

**Figure 2.**
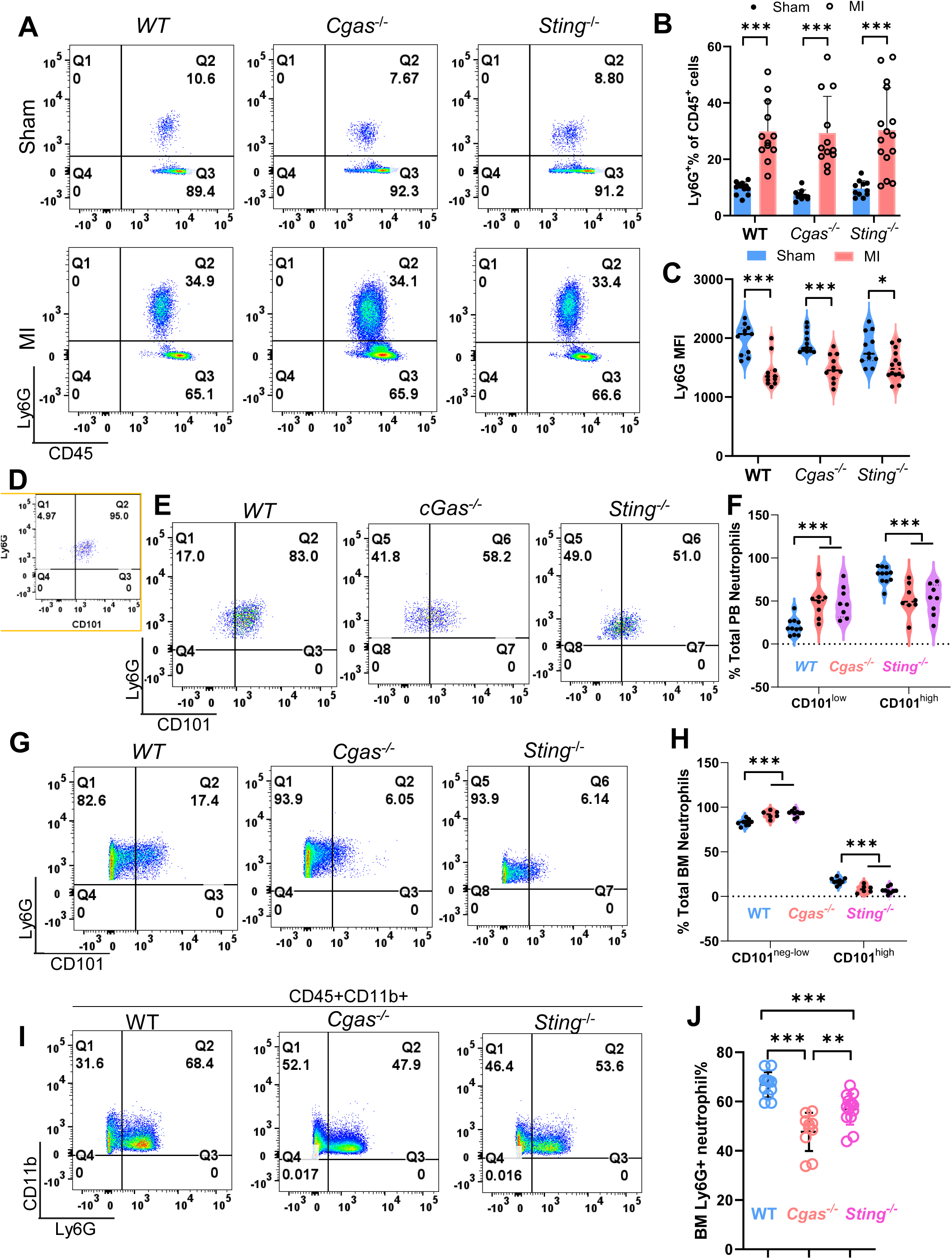
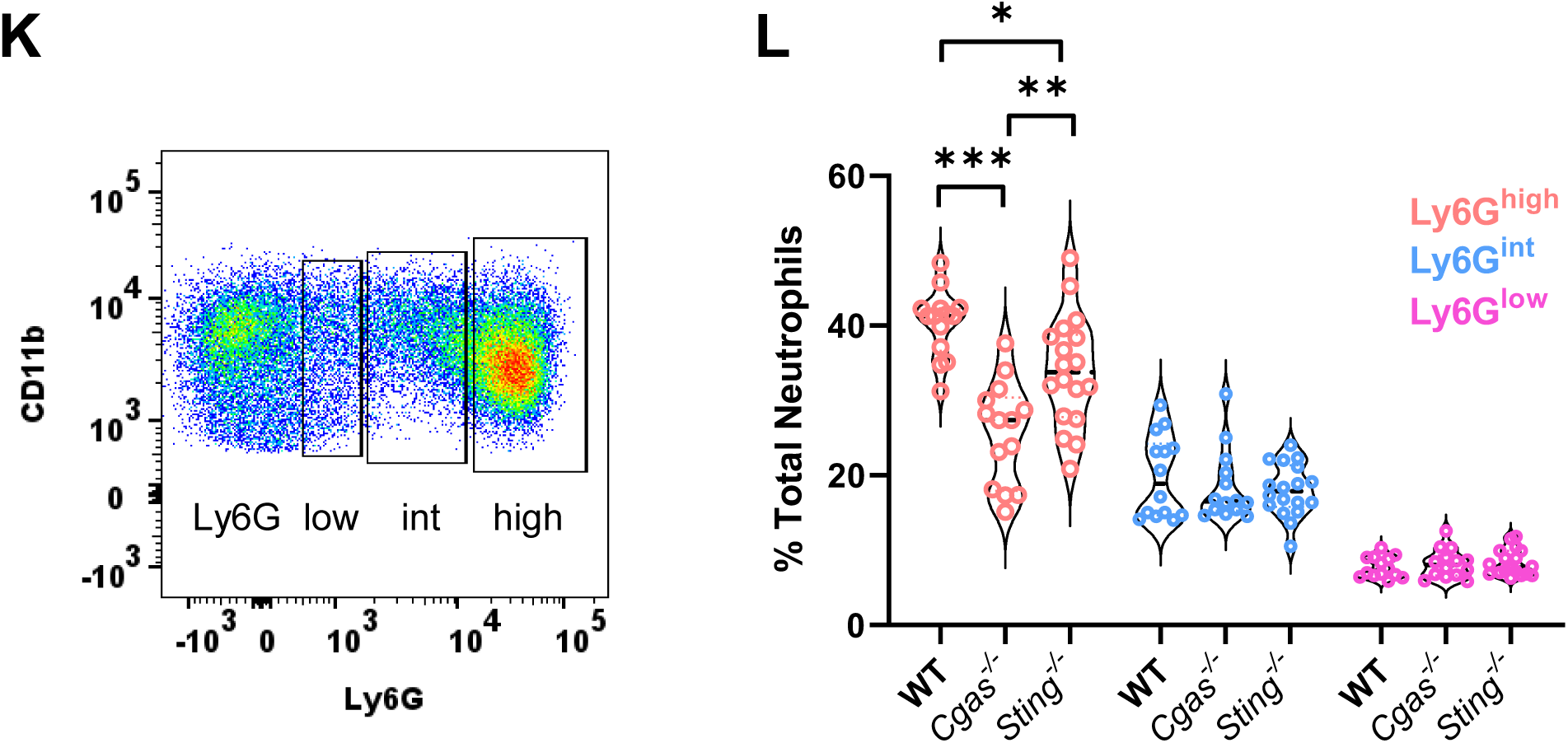
Altered neutrophil phenotype post-MI in *Cgas*- and *Sting*-deficient mice. Mice from the indicated genotype underwent sham- or MI-procedure. Three days after the procedure, peripheral blood (PB, **A**-**C**) or bone marrow (BM, **G**-**L**) was analyzed. **A**-**C**. PB neutrophils were identified by flow cytometry as the CD45^+^CD11b^+^Ly6G^+^ population. Representative scatter plot (**A**) and summary data for percent neutrophils to the total leukocyte (CD45^+^) (**B**) and Ly6G MFI (**C**) are shown. **D**. Gating parameter setup using WT sham PB to differentiate CD101^low^ and CD101^high^ neutrophil population. The proportion of CD101^low^ and CD101^high^ neutrophils from the PB (**E**-**F**) and BM (**G**-**H**) after MI. **I**-**J**. Representative scatter plot (**I**) and summary data (**J**) of the entire neutrophil compartment in BM after MI determined as CD45^+^CD11b^+^ with any level of Ly6G expression. **K**-**L**. The neutrophil compartment, as determined in J, was grouped based on Ly6G expression (K), and each group’s proportion of the total neutrophil population was determined (**L**). *p<0.05. **p<0.01. ***p<0.001.

We assessed the maturity of circulating neutrophils. Firstly, we measured the median fluorescent intensity (MFI) of Ly6G because this marker progressively increases from the early lineage-committed progenitors to mature neutrophils. Three days after myocardial infarction (MI), PB neutrophils showed a significant decrease in Ly6G MFI compared to sham controls (**Figure 2C**), suggesting earlier release from the bone marrow at less mature stages of differentiation. This decrease was consistent across WT, *Cgas*^-/-^, and *Stin*g^-/-^ PB neutrophils. We next evaluated CD101 in neutrophils produced by the three genotypes of mice after MI. CD101 is a marker that distinguishes proliferative neutrophils (CD101^-^) from terminally differentiated neutrophils (CD101^+^)^10^. In sham WT mice, 95% of PB neutrophils were CD101^high^ (**Figure 2D**), a proportion similar in *Cgas*^-/-^, and *Stin*g^-/-^ neutrophils (data not shown). Using the gating parameter set in **Figure 2D**, CD101^high^ PB neutrophils decreased to 80% in WT neutrophils and 50% in *Cgas*^-/-^ and *Stin*g^-/-^ counterparts post-MI (**Figure 2E, 2F**). In contrast to the PB, BM neutrophils after MI were predominantly the CD101^neg-low^ population, accounting for 80% of WT neutrophils and significantly more in *Cgas*^-/-^ and *Stin*g^-/-^ mice (**Figure 2G, 2H**). CD101 has been utilized as a marker of neutrophil maturation in recent studies^10, 27–29^. Our findings indicate that it differentiated the maturation states of neutrophils produced by the three types of mice after myocardial infarction.

The higher percentage of CD101^neg-low^ neutrophils in *Cgas*- and *Sting*-deficient mice from both PB and BM implies a role for the cGAS-STING pathway in neutrophil differentiation and maturation. To explore this, we characterized the total BM neutrophil compartment after MI. BM neutrophils were defined as CD45^+^CD11b^+^ and Ly6G^low-to-high^ to include cells from very early differentiation stages (Ly6G^low^) to the final population containing band and segmented cells (Ly6G^high^). As expected, neutrophils were a major cell type produced in the BM after MI, accounting for 68% of total myeloid cells in WT mice (**Figure 2I**). Compared to WT BM, BM from *Cgas*^-/-^ and *Sting^-^*^/-^ mice, especially *Cgas*^-/-^, had lower percentages of neutrophils after MI (**Figure 2I, 2J**). We divided all Ly6G^+^ BM neutrophils into three subpopulations: Ly6G^low^, Ly6G^int^, and Ly6G^high^, correlating with pre-neutrophils (PN), immature neutrophils (IN), and band and segmented cells (BC/SC), respectively^10, 11^ (**Figure 2K**). Interestingly, the percentage of Ly6G^high^ neutrophils was significantly reduced in *Cgas*^-/-^ and *Sting^-^*^/-^ mice (**Figure 2L**), especially the *Cgas-* deficient mice. In aggregate, these data suggest that although *Cgas*^-/-^ and *Sting*^-/-^ mice mount a similar neutrophilic response to MI, indicating that neutrophil mobilization from BM to PB is preserved, the cGAS-STING pathway promotes neutrophil differentiation and maturation during ischemia-induced neutropoiesis.

### Acute ischemia promotes pathways essential to neutrophil effector function in bone marrow

We performed transcriptome analysis using RNA-Seq to investigate how BM neutrophils respond to MI and how the cGAS-STING pathway influences this response. We first purified neutrophils (CD45^+^CD11b^+^CCR2^-^Ly6C^int^Ly6G^+^) using FACS (**Figure 3A**) from BM three days after MI or sham procedure in WT mice. WT BM neutrophils after MI had 862 upregulated and 194 down-regulated genes (**Figure 3B**). Down-regulated transcripts included transcription factors (TFs) such as the Fos family members (*Fos, Fosb, and Fosl1*) and *Egr1* (**Figure 3B**). These TFs are highly expressed in mature neutrophils at homeostasis and increase transcriptional activity during normal neutrophil differentiation^10, 11, 30^. However, during emergent neutropoiesis induced by an infectious agent’s challenge, these TFs’ expression and their controlled transcriptional programs were suppressed in mature neutrophils^11^. Our findings here align with previous reports on the dynamics of these TFs following inflammatory stimuli.

**Figure 3.**
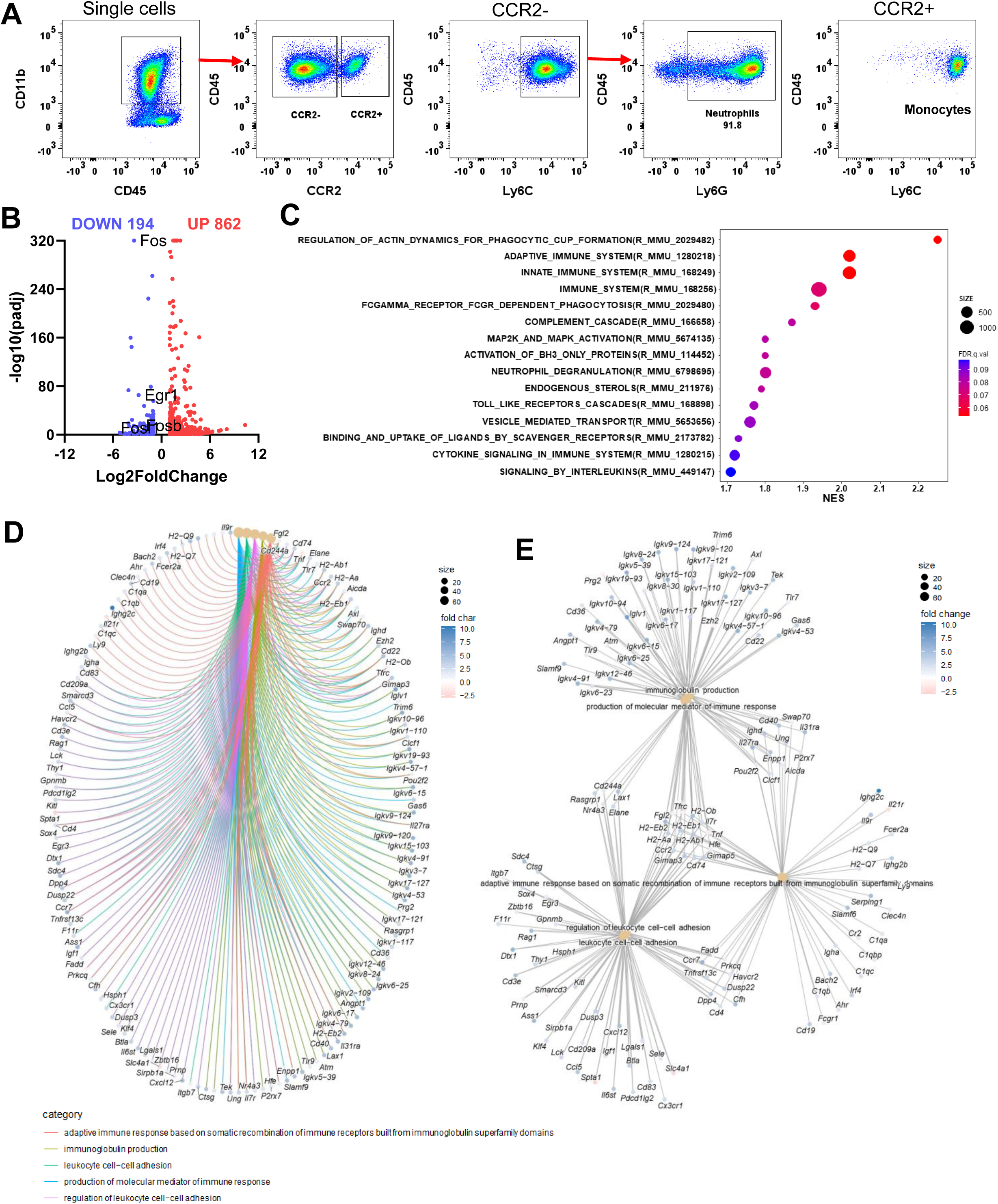
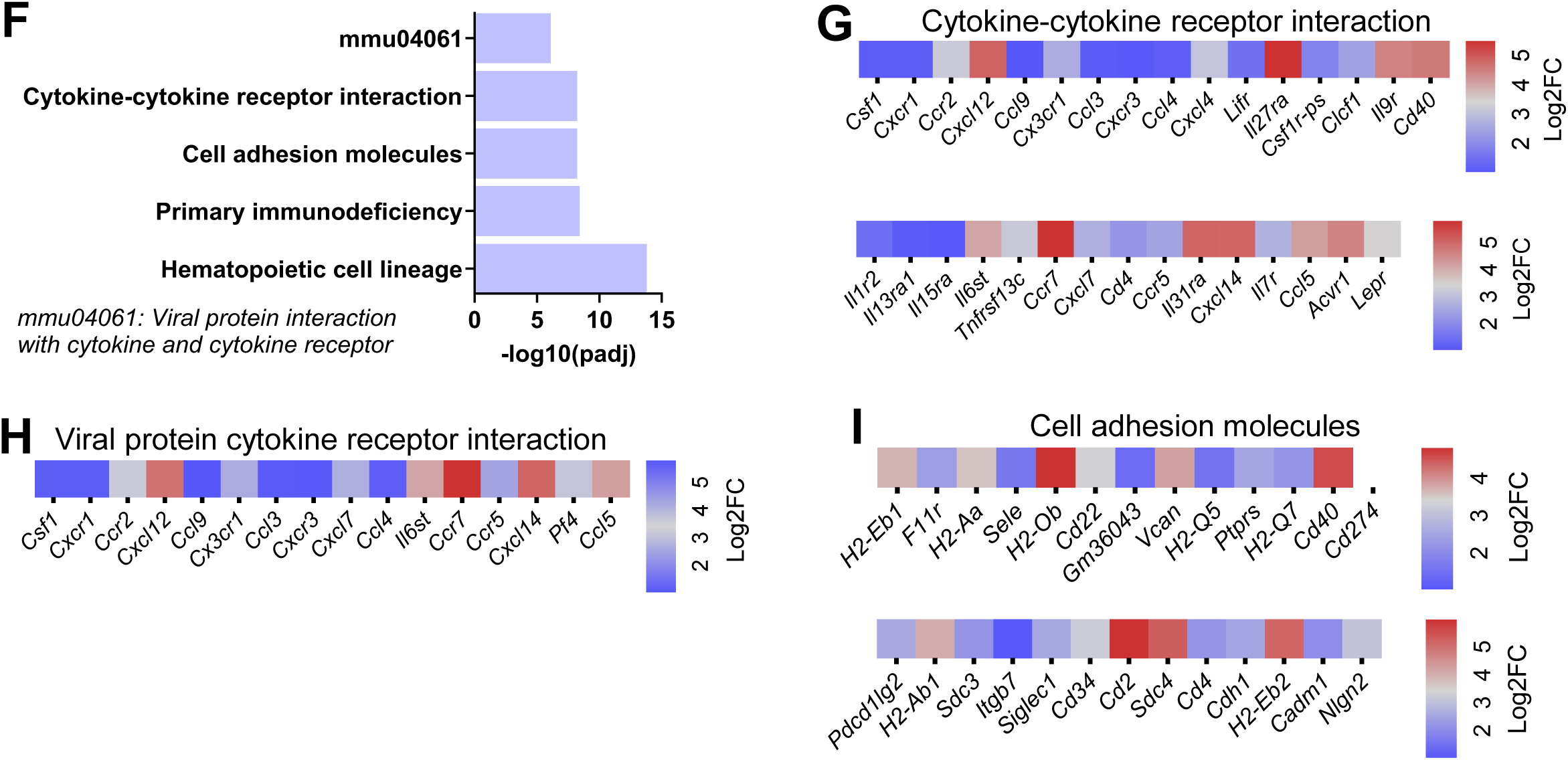
Acute ischemia promotes functionality of neutrophils in the bone marrow. BM neutrophils purified by FACS were used to extract total RNA for RNA-sequencing. Transcriptomic changes in WT BM neutrophils three days after MI-procedure was compared to the neutrophils from the BM of sham-operated mice. **A**. FACS gating strategy to purify the CD45^+^CD11b^+^CCR2^-^ Ly6C^int^Ly6G^+^ neutrophil population. **B**. Volcano plot of differentially expressed genes using cutoffs of Log2 fold change (Log2FC) equal to or greater than 1 and adjusted p (padj) value <0.05. **C**. Normalized enrichment score of the top 15 up-regulated processes determined by Reactome analysis. **D**. Cnetplot of indicated GO processes and specific genes that were differentially regulated. **E.** Gene network analysis. Nodes represent GO processes; edges link the nodes to specific genes. **F**. The most up-regulated pathways in MI neutrophils compared to sham neutrophils determined by KEGG analysis. **G**-**I**. Heatmaps to visualize differentially up-regulated genes by ischemia in three of the most enriched pathways from KEGG analysis.

Next, we performed pathway enrichment analysis. **Figure 3C** illustrates the Reactome processes enriched in BM neutrophils by MI, including phagocytosis, degranulation, vesicle trafficking, cytokine signaling, and TLR signaling. These pathways are essential to neutrophil migration, engulfing tissue debris, releasing enzymes stored in granules, and responding to cytokines. Additionally, MAP kinase activation, which promotes survival, and BH3 activation, which triggers apoptosis, were among the most enriched processes (**Figure 3C**). This suggests that ischemia regulates the neutrophil life cycle starting in the BM, indicating that neutrophil life expectancy is tightly controlled to minimize damage from these potent cells.

**Figure 3D** is a cnetplot integrating the differentially regulated genes and the fold changes for the most enriched GO (Gene Ontology) terms in BM neutrophils after MI compared to sham-operated mice. The size of the gene pools for each GO term was depicted. The cnetplot also revealed genes that belonged to multiple biological processes. **Figure 3E** is a gene network analysis, demonstrating clusters of genes shared among the enriched processes. The core enriched GOs (**Figure 3D, 3E**) included pathways involved in generating molecular mediators of the immune response (*Elane*, *Tnf*, *Ccl2*, *Tlr7*, *Tlr9*, *Tfrc*) and mediating leukocyte cell adhesion (*Cxcl12, Cx3cr1, Klf4, Itgb7)*. Interestingly, processes related to immunoglobulin production and antigen processing and presentation were enriched. The latter was indicated by the upregulation of histocompatibility 2 (H2) isotypes, which are components of the MHC class II complex.

The KEGG analysis identified the hematopoietic cell lineage as a top upregulated pathway by ischemia (**Figure 3F**), consistent with the enhanced hematopoietic process triggered by MI. To visualize the specific genes that were differentially regulated in the KEGG pathways, we plotted heatmaps of significantly upregulated genes by ischemia. In the "Cytokine-cytokine receptor interaction" pathway, chemokine receptors, including *Ccr2/5/7*, *Cxcr1/3*, and the receptor that binds IL1A and IL1B were the most upregulated genes (**Figure 3G**). Ischemia also promoted the expression of genes in the "Viral protein interaction with cytokine and cytokine receptor" pathway, including members from the *Ccl-Ccr, Cxcl-Cxcr,* and *Cx3cl-Cx3cR* cytokine-receptor families (**Figure 3H**). This pathway comprises targets of viral proteins that subvert and modulate the host immune defense system, which explains the overlap with the cytokine-cytokine receptor pathway (**Figure 3G**). Additionally, **Figure 3I** depicts transcripts from the "Cell adhesion molecules" pathway induced in BM neutrophils after acute MI. The upregulation of seven components of the major histocompatibility complex (MHC) II, including *H2-Eb1, H2-Aa, H2-Ob, H2-Q5, H2-Q7, H2-Ab1, H2-Eb2,* and *Pd-l1/Cd274* suggests that MI promotes an antigen-presenting function during neutrophil development in the BM, consistent with the enrichment analysis presented in the cnetplots (**Figure 3D, 3E**). The role of neutrophils as antigen-presenting cells has been proposed by multiple recent studies^31–34^, but such a role in MI is unknown.

In summary, transcriptome analysis of BM neutrophils after MI showed upregulation of processes essential to neutrophil function, including phagocytosis, degranulation, cytokine production, cytokine-receptor interaction, and cell adhesion. These data indicate that ischemia enhances neutrophil functionality before their release into the circulation and migration to the infarct tissue, suggesting that altering the neutrophil cell state is part of the neutropoiesis triggered by MI in addition to the neutrophilic response (increase in neutrophil numbers). Interestingly, acute ischemia also activated pathways related to adaptive immunity. While the functional relevance of these findings remains unclear, they warrant further investigation, as they may indicate that neutrophils play a role in modulating adaptive immune responses in the infarcted heart.

### cGAS-STING pathway regulates the transcriptional response of BM neutrophils after MI

To determine if cGAS and STING regulate the differentiation and maturation of neutrophils after MI, we performed transcriptome analysis using RNA-Seq on BM neutrophils from *WT*, *cGas*^-/-^ or *Sting*^-/-^ mice three days post-MI. Neutrophils were sorted by FACS using the gating strategy outlined in **Figure 3A**. Principal component analysis (PCA) (**Figure 4A**) revealed a clear separation of the transcriptomes from the three genotypes. Compared to their WT counterparts, *Cgas*-deficient BM neutrophils had 1016 down-regulated and 772 upregulated genes after MI, using cut-off criteria of Log2 fold change (Log2FC) ≥ 1 and adjusted p-value < 0.05 (**Figure 4B**). *Sting*-deficient neutrophils showed 460 down-regulated and 702 upregulated genes (**Figure 4C**). We first performed a cnetplot analysis of the *Cgas*^-/-^ BM neutrophils compared to WT neutrophils after MI. **Figure 4D** depicts the five core processes down-regulated by *Cgas* deletion. Among them are the activation of immune responses, cellular response to IFNB, and defense response to viruses. In **Figure 4E**, the nodes represent the different processes, and the edges connect the process to differentially regulated genes. **Figure 4E** also visualizes the distribution of common and distinct genes among nodes. Next, we depicted the ten most enriched GO terms in down-regulated genes due to *Sting* deletion in **Figure 4F**, which included six pathways involving virus defense, and cellular response to IFNB, significantly overlapping with those of *Cgas*^-/-^ neutrophils (**Figure 4D, 4E**). The top five enriched KEGG pathways for down-regulated transcriptomes in *Cgas*^-/-^ (**Figure 4G**) and *Sting*^-/-^ (**Figure 4H**) BM neutrophils included shared pathways such as "Viral protein interaction with cytokine and cytokine receptor" and "Cell adhesion molecules," as well as unique pathways. Because the shared processes were the top upregulated pathways by ischemia (**Figure 3F**), the data suggests that the cGAS-STING is an essential pathway to enhance the functionalities of neutrophils during MI-triggered neutropoiesis. Furthermore, KEGG enrichment analysis demonstrated that the "Nod-like receptor signaling pathway" was enriched in the down-regulated transcriptomes in both *Cgas*^-/-^ (ranked No.10) and *Sting*^-/-^ (ranked No.1) neutrophils compared to the WT counterparts. **Figure 4I** is the heatmap depicting specific genes in "Nod-like receptor signaling pathway" down-regulated by *Cgas* or *Sting* deletion. Interestingly, many members of this pathway, such as *Irf7, Oas1a, Oas1b, Oas1g, Oas2, Ifi204, Gbp3, and Gbp5*, are also ISGs controlled by the cGAS-STING, suggesting a connection between IFN signaling and the Nod-like receptor pathway. Prior studies showed that the Nod-like receptor family (e.g., NLRP3) is necessary for neutrophil generation after MI^35, 36^. Loss of cGAS and STING could impair Nod signaling, thereby dampening neutrophil production after MI (**Figure** 2I to 2L).

**Figure 4.**
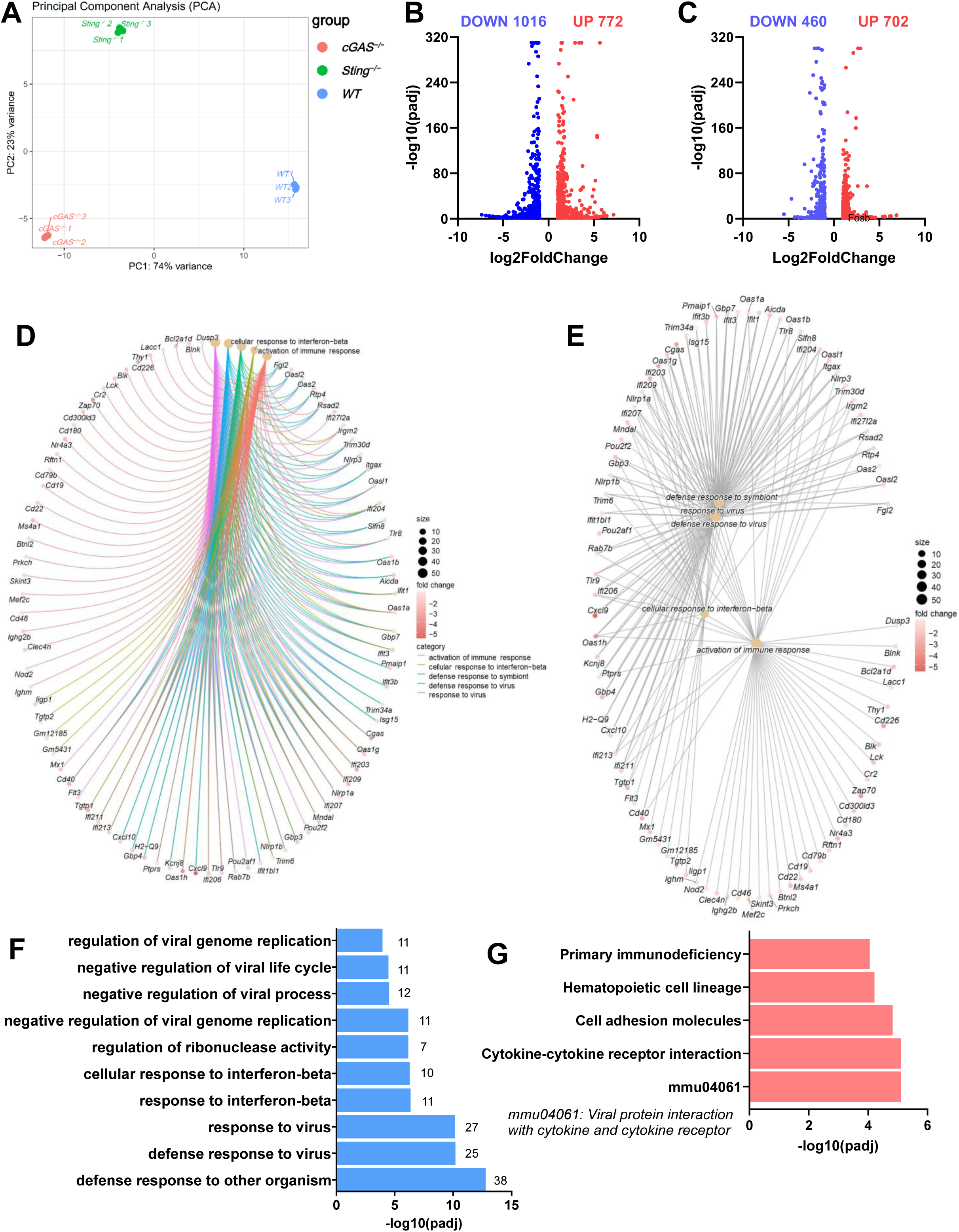
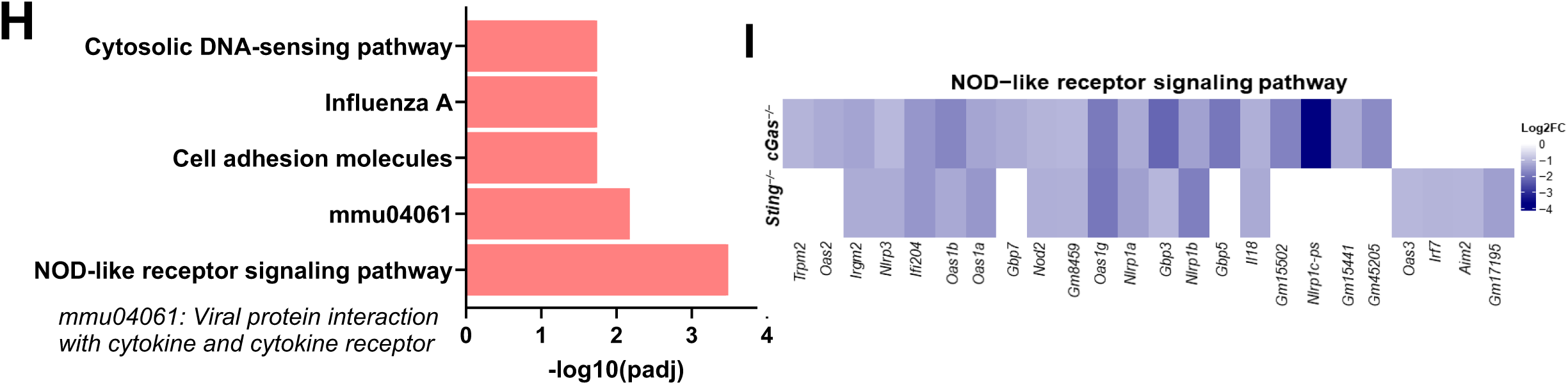
cGAS-STING pathway promotes neutrophil functionality during neutropoiesis after MI. Neutrophils were isolated from the BM of WT, *Cgas*^-/-^, and *Sting*^-/-^ mice three days after MI using the gating strategy described in Figure 3A. **A**. PCA plot of the RNA-sequencing analysis. **B**-**C**. Volcano plots of differentially expressed genes in *Cgas*^-/-^ neutrophils compared to WT neutrophils (**B**) and *Sting*^-/-^ neutrophils compared to WT neutrophils (**C**). **D**. cnetplot of indicated GO processes and specific genes that were differentially regulated in *Cgas*^-/-^ BM neutrophils compared to WT BM neutrophils after MI. **E**. Gene network analysis of *Cgas*^-/-^ BM neutrophils compared to WT neutrophils. **F**. The top 10 most enriched GO terms in the down-regulated genes in *Sting*^-/-^ neutrophils compared to WT neutrophils. **G**-**H**. The top five enriched KEGG pathways in the down-regulated transcriptomes in *Cgas*^-/-^ (**G**) and *Sting*^-/-^ (**H**) neutrophils compared to WT neutrophils. **I**. The heatmap demonstrates the significantly down-regulated genes from the “Nod-like receptor signaling pathway” of KEGG analysis in neutrophils of indicated genotypes compared to WT neutrophils.

We compared the cGAS- and STING-regulated transcriptomic changes after MI by pairwise comparing *cGas*^-/-^ or *Sting*^-/-^ BM neutrophils to wild-type (WT) counterparts. To analyze gene co-expression and anti-correlation patterns in neutrophils from the three genotypes, we performed hierarchical clustering and generated a heatmap (**Figure 5A**). This analysis identified four clusters of genes with similar expression patterns, separated by genotypes. We performed GO enrichment analyses on these clusters. Clusters 2 and 4 will not be discussed further as they involved limited GO terms. For example, Cluster 4 included four hemoglobin subunit genes, *Snora31* (small nucleolar RNA, H/ACA box 31), and *Ear1* (eosinophil-associated ribonuclease 1). The genes in Cluster 1 were predominantly upregulated in WT BM neutrophils and down-regulated in both *Cgas^-/-^* and *Sting*^-/-^ BM neutrophils after MI. The GO terms enriched in this cluster included transcriptomic programs for immune response activation, immune signaling pathways, leukocyte activation, cytokine production, and phagocytosis (**Figure 5B**). The enriched processes of Cluster 1 further support the analysis presented in **Figures 4D** to **4F** that inhibiting cGAS-STING dampens the neutrophil priming triggered by MI.

**Figure 5.**
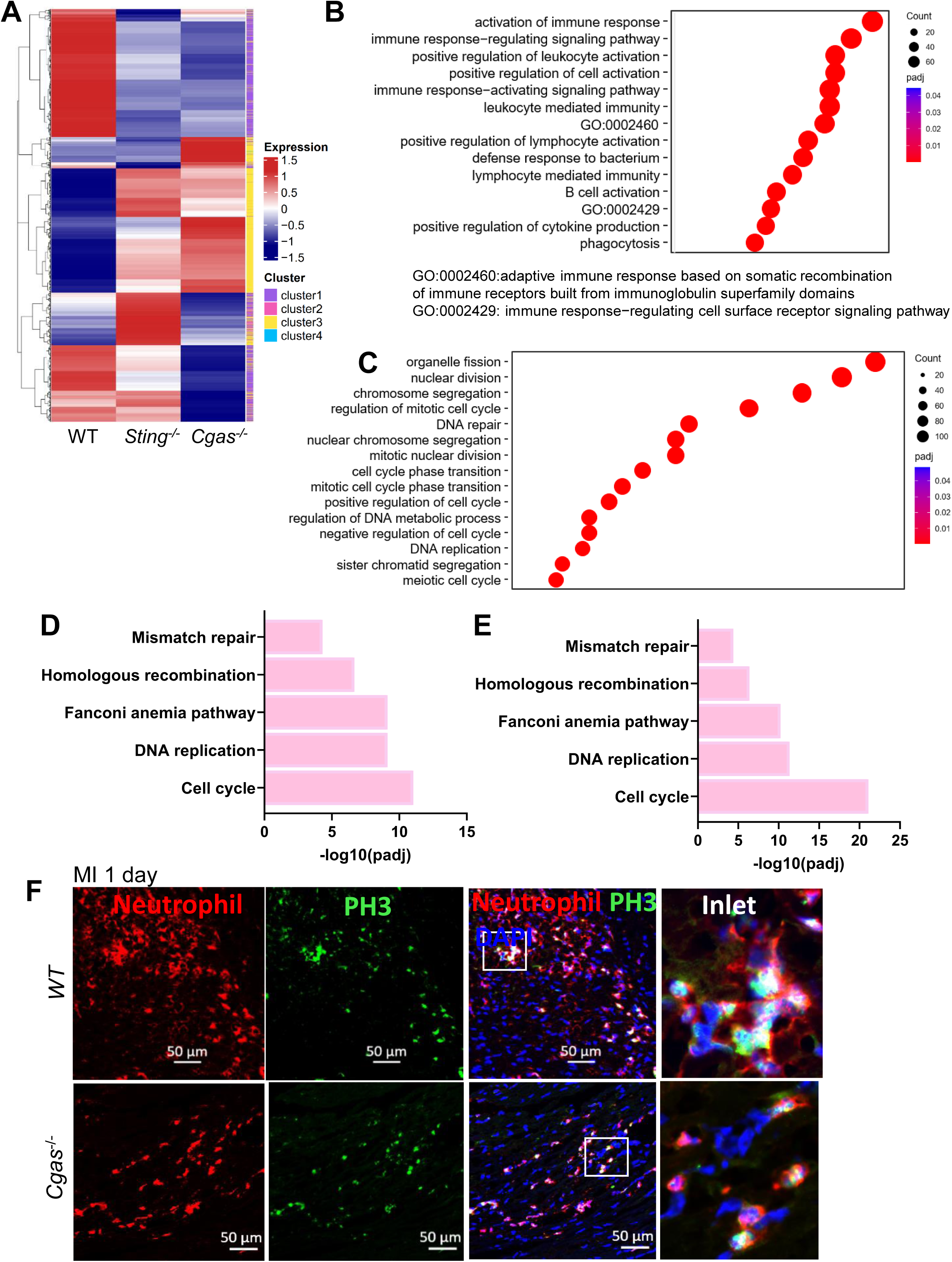
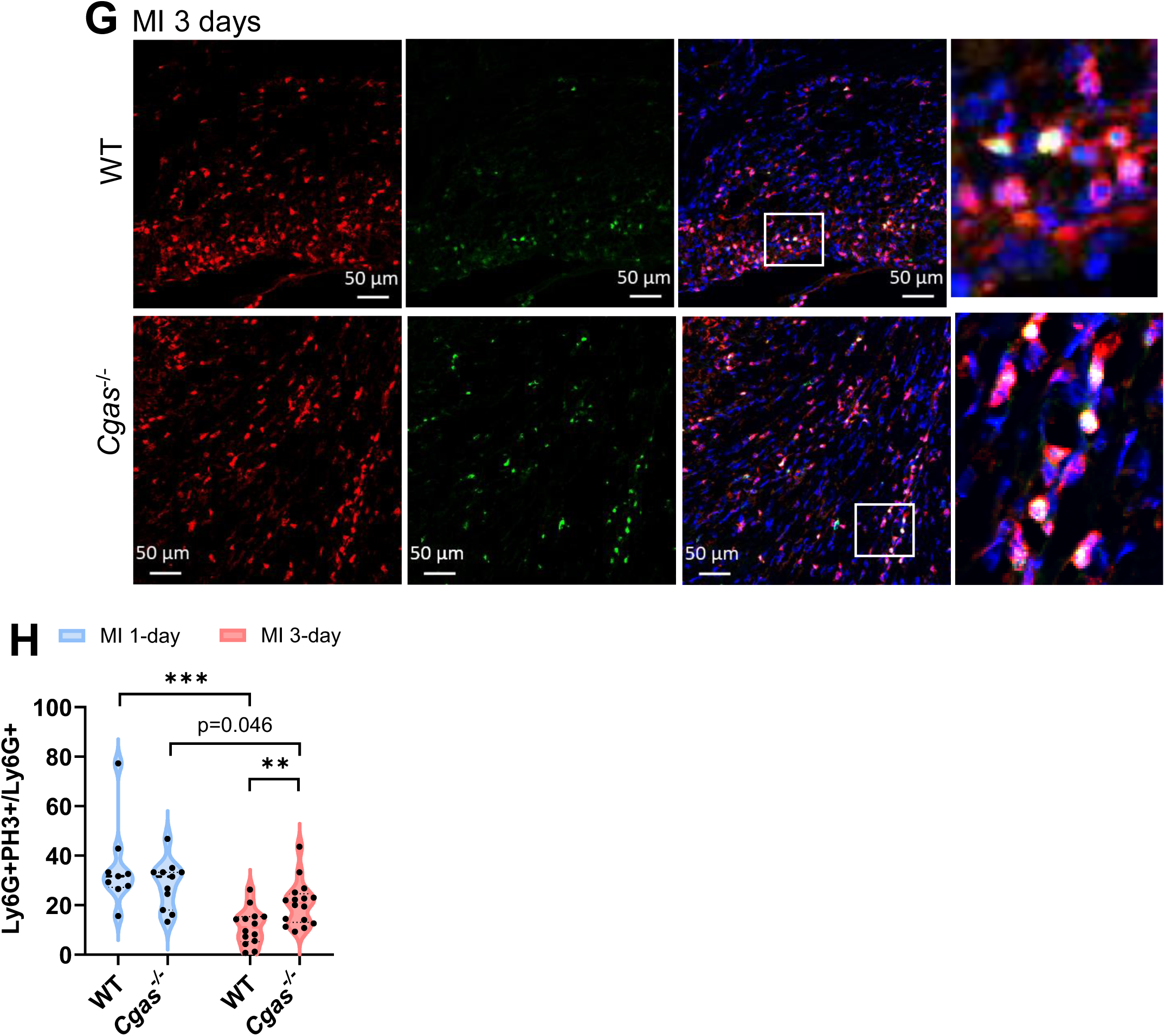
Augmented cell cycle programs in *Cgas*- and *Sting*-deficient neutrophils after MI. **A**. Hierarchical heatmap of the RNA-sequencing data to identify gene expression patterns. Four sub-clusters were identified. **B**-**C**. GO term enrichment analysis of sub-cluster 1 (**B**) and sub-cluster 3 (**C**). **D**-**E**. Enriched KEGG pathways from the up-regulated genes of *Cgas*^-/-^ BM neutrophils (**D**) or *Sting*^-/-^ BM neutrophils (**E**) compared to WT BM neutrophils after MI. **F**-**G**. Immunohistochemistry of cryosections of ischemic hearts prepared either 24 hours (**F**) or 3 days (**G**) after MI surgery. **H**. Images acquired as in **F**-**G** were used to count the percentage of PH3+neutrophils to total neutrophils. **p<0.01. ***p<0.001.

We therefore focused on Cluster 3, which contained genes uniquely upregulated in *Cgas*^-/-^ and *Sting*^-/-^ BM neutrophils after MI but down-regulated in WT counterparts. This cluster’s most enriched GO terms were nuclear division, chromosome segregation, cell cycle, mitosis, and DNA replication (**Figure 5C**). KEGG analysis of the upregulated transcriptomes revealed the top 5 pathways identical in *Cgas*^-/-^ (**Figure 5D**) and *Sting*^-/-^ (**Figure 5E**) neutrophils, including cell cycle, DNA replication, and DNA repair. The upregulated cell cycle programs in *Cgas*- and *Sting*-deficient neutrophils were unexpected. To validate these findings at the protein level, we stained cardiac neutrophils with the proliferation marker phospho-histone 3 (PH3) (**Figure 5F and 5G**). One day after MI, more than one-third of neutrophils in the infarct region were positive for PH3 in both *WT* and *cGas*^-/-^ mice (**Figure 5F** and **5H)**. PH3+ neutrophils decreased significantly in WT three days after MI but were maintained in the *Cgas*^-/-^ MI hearts (**Figure 5G** and **5H**). These results align with a recent proteomics study that showed BM neutrophil cell cycle proteins were the most prominently increased during emergent neutropoiesis and remained elevated after migrating into peripheral blood and tissues after systemic infection^37^.

Neutrophils rapidly lose proliferation capacity after lineage commitment, and only a small percentage of Ly6G^low^ cells can divide^10^. The neutrophil population used in the RNA-seq analysis (**Figure 3A**) consisted of largely non-proliferating cells. PH3+ neutrophils in the infarct tissue, where typically only terminally differentiated neutrophils are present, suggest a dissociation between cell cycle-related programs and neutrophil proliferation. This dissociation is more prominent in *Cgas*- and *Sting*-deficient neutrophils.

### cGAS-STING pathway is essential for ISG expression in neutrophils after MI

Single-cell studies have identified a subset of neutrophils with a transcriptional (mRNA) signature of IFN-ISGs at homeostasis that expand during emergent neutropoiesis triggered by infection^11, 19^. To determine if the cGAS-STING pathway regulates ISG expression in neutrophils, we selected 62 ISGs based on literature and database searches^11, 17, 19, 38^ and analyzed their expression pattern three days after MI in neutrophils from BM (**Figure 6A**) or infarct tissue (**Figure 6B**). Most ISGs (83-90%) were downregulated in *Cgas-* or *Sting*-deficient BM neutrophils after MI compared to their WT BM counterparts (**Figure 6A**), with 70% and 55% of the downregulated ISGs showing more than a 2-fold change (Log2FC≥1) and an adjusted p-value < 0.05, respectively. No ISGs expressed at lower levels in WT compared to *Cgas*^-/-^ and *Sting*^-/-^ neutrophils met these criteria.

**Figure 6.**
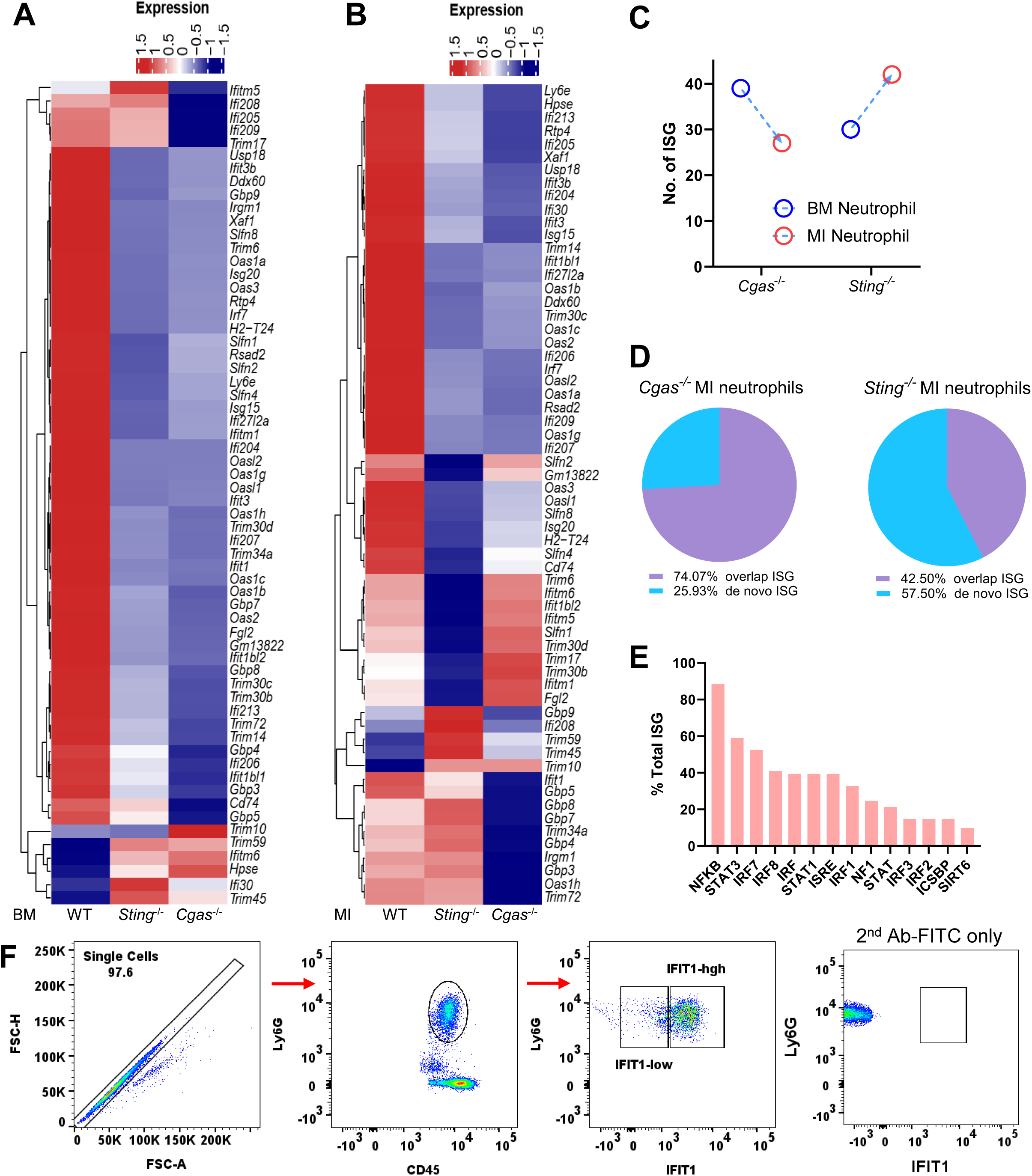
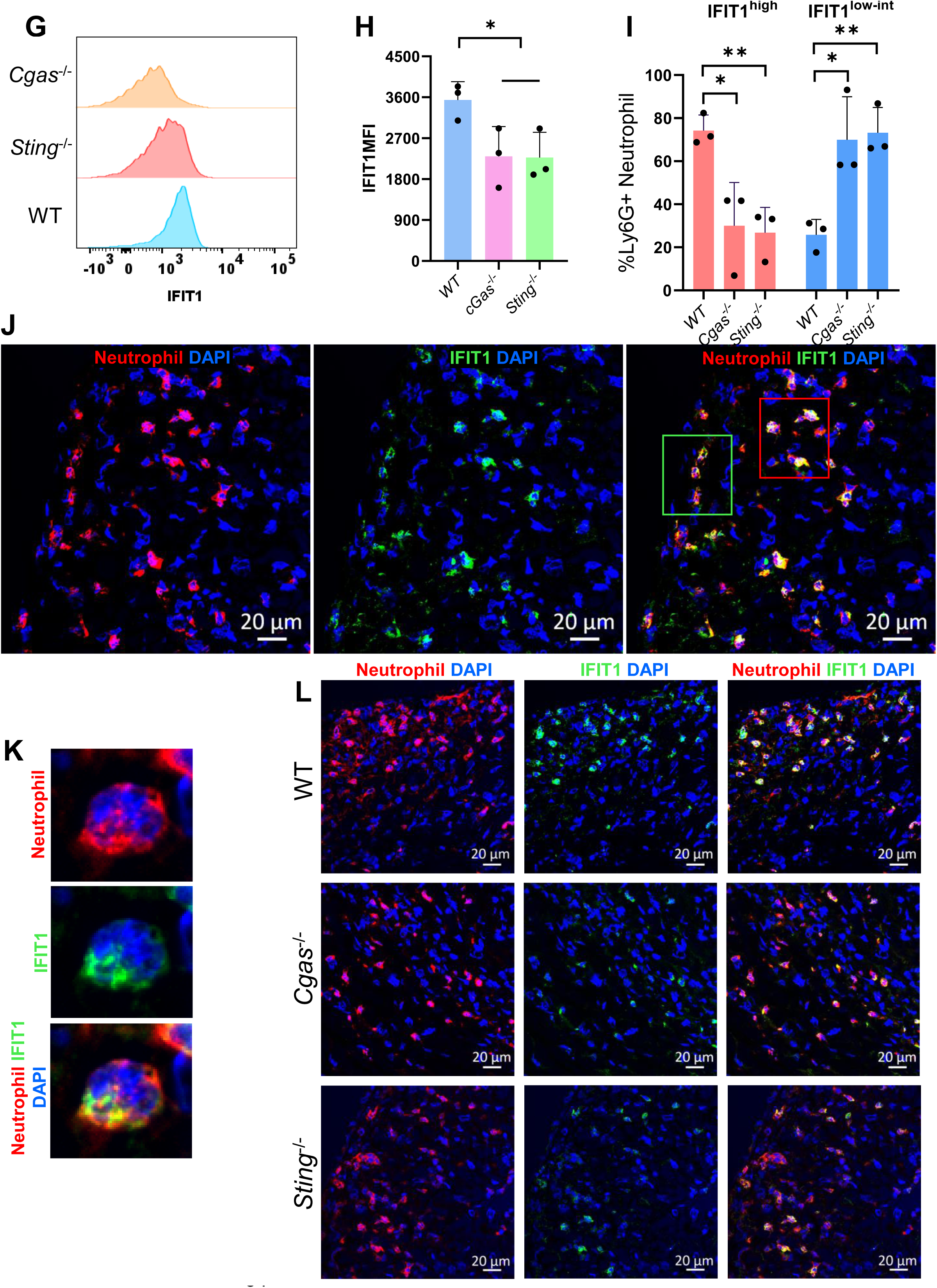
cGAS-STING pathway promotes the ISG signature in neutrophils. **A**-**E**. The expression pattern of 62 ISGs was analyzed from the RNA-seq data generated from BM or cardiac neutrophils after MI. **A**. Hierarchical heatmap of the 62 ISGs in BM neutrophils after MI. **B**. Neutrophils were isolated from ischemic tissue using Ly6G-beads positive selection three days after MI. Total RNAs were extracted from the purified MI neutrophils followed by RNA sequencing. The hierarchical heatmap for the same 62 ISGs (as in **A**) is shown. **C**. The difference in the number of significantly down-regulated ISGs between BM *Cgas^-^*^/-^ and MI *Cgas^-^*^/-^ neutrophils, and that between the BM *Sting*^-/-^ and MI *Sting*^-/-^ neutrophils. **D**. The percentage of ISGs induced by the ischemic environment (de novo) and overlap ISGs (down-regulation in both BM and MI tissue) in *Cgas^-^*^/-^ and *Sting*^-/-^ neutrophils. **E.** The percentage of the 62 ISGs that have binding sites for the indicated transcription factors. **F**-**I**. PB neutrophils three days after MI were stained for IFIT1. Gating strategy to differentiate the IFIT1^low^ and IFIT1^high^ neutrophils (**F**). Representative histogram of IFIT1+ neutrophil distribution (**G**) and summary data for IFIT1 MFI (**H**). The percentage of neutrophils that are IFIT1^low^ and IFIT1^high^ (**I**). **J**-**L**. IFIT1 immunostaining of cardiac sections prepared three days after MI. IFIT1+ cells are exclusively neutrophils (**J**). The distinct pattern of IFIT1 and anti-neutrophil surface antigen staining (**K**). The IFIT1+ neutrophils in WT, *Cgas*^-/-^, and *Sting*^-/-^ MI hearts (**L**). Images are representative of 3 to 4 MI hearts. *p<0.05. **p<0.01.

In neutrophils isolated from infarct tissue at the same time point after MI, ISG expression followed a similar pattern to that in BM neutrophils, with the majority of ISGs (79%) decreased in *Cags*^-/-^ or *Sting*^-/-^ neutrophils. Of these, 45-65% met the strict criteria mentioned above (**Figure 6B**). The only ISG with lower expression in WT MI neutrophils that met the selection criteria (log2FC≥1 and padj<0.05) was *Ifi208.* The total number of significantly downregulated ISGs decreased in *Cgas*^-/-^ MI neutrophils compared to *Cgas*^-/-^ BM neutrophils (27 vs. 39, respectively). Conversely, the number of downregulated ISGs was higher in *Sting*^-/-^ MI neutrophils compared to the *Sting*^-/-^ BM cells (41 vs. 30) (**Figure 6C**). Downregulated ISGs in *Cgas*^-/-^ MI neutrophils largely overlapped with those in their BM counterparts, whereas 50% of the significantly downregulated ISGs in *Sting*^-/-^ MI neutrophils were specifically induced by the ischemic environment ("de novo") (**Figure 6D**). These "de novo" differentially expressed ISGs in MI neutrophils were comparable between WT and *Sting*^-/-^ BM neutrophils. The overwhelming down-regulation of ISG with cGAS and STING inhibition highlights the essential role of this pathway in controlling ISG expressions in neutrophils.

The number of ISGs differentially regulated by cGAS and STING was considerably larger than those reported to be regulated by IRF3^19^. One likely explanation for this difference is that ISGs are regulated by multiple transcription factors (TFs). Using bioinformatic analysis, we identified binding sites for multiple TFs in this group of 62 ISGs, including NFKB, STAT3, IRF7, IRF8, STAT1, ISRE, IRF1, IRF2, IRF3, NF1, ICSBP, and SIRT6 (**Figure 6E).** 89% of the ISGs had at least one binding site for NFKB, followed by STAT3 (59%) and IRF7 (53%) (**Figure 6E**). IRF3 binding sites were detected in 15% of the ISGs (**Figure 6E**). NFKB, IRF3, and IRF7 are all downstream targets of cGAS-STING activation, suggesting that the broader scope of ISG regulation may be due to NFKB and IRF3 activation by cGAS-STING. The robustness of ISG expression likely also depends on the availability of these TFs in the different milieus where neutrophils reside.

In summary, our data suggest that cGAS-STING promotes ISG expression during neutrophil differentiation and in response to MI injury. The data also indicate that STING may play a broader role in IFN-activated programs, especially in response to the MI environment.

Most published data regarding neutrophils with ISG signatures were performed at the transcriptional level. To assess ISG protein expression in neutrophils and define the role of cGAS-STING, we assess IFIT1^+^ neutrophils by flow cytometry after intracellular staining three days after MI. We used neutrophils from peripheral blood as they more reliably reflect the overall generation of ISG-expressing neutrophils, and prior work suggested IFIT1^+^ neutrophils were mature neutrophils existing mainly in peripheral blood and spleen^11^. **Figure 6F** shows the scatter plots of the gating sequence for IFIT1^+^ neutrophils and the control performed using only the FITC-conjugated secondary antibody. **Figure 6G** depicts the neutrophil distribution based on **the** IFIT1 signal by flow cytometry. The mean fluorescence intensity (MFI) of IFIT1 (**Figure 6H**) was decreased in *Cgas^-/-^* and *Sting*^-/-^ neutrophils, suggesting these cells express less IFIT1 protein. In addition, the percentage of IFIT1^high^ neutrophils was significantly smaller in *Cgas^-/-^* and *Sting*^-/-^ neutrophils compared to WT (**Figure 6I**).

To assess the IFIT1 neutrophil population in the infarct tissue, we co-labeled heart cryosections with anti-neutrophil (red) and IFIT1 (green) antibodies. Interestingly and surprisingly, IFIT1^+^ cells were almost exclusively neutrophils (**Figure 6J**). Among the neutrophils, some cells stained strongly with IFIT1 (red inlet, **Figure 6J**), while others exhibited a weaker IFIT1 signal (green inlet, **Figure 6J**). Very few neutrophils were IFIT1 negative. These data suggest non-uniform IFIT1 expression in MI neutrophils. The IFIT1 staining pattern is distinct from the anti-neutrophil staining targeting Ly6B.2, a GPI-anchored cell surface protein (**Figure 6K**). Next, we assessed the IFIT1^+^ neutrophil population in the infarct hearts from *Cgas*- and *Sting*-deficient mice. Neutrophils from both knockout mice showed reduced IFIT1 expression (**Figure 6L**) but not completely absent. These data suggest a crucial role for the cGAS-STING pathway in regulating IFIT1 expression in neutrophils, which is in line with the RNA-seq analysis of ISGs.

### cGAS-STING promotes neutrophil function in the ischemic myocardium

Transcriptional profiling of BM neutrophils after MI revealed upregulation of canonical functions even before neutrophils reach the infarct tissue. The loss of cGAS-STING attenuated this upregulation in the BM. To determine if the transcriptomic changes observed in the BM were maintained in neutrophils that had reached the ischemic environment, we purified neutrophils from infarct tissue three days after MI using Ly6G UltraPure Microbeads^39, 40^ and performed RNA-seq analysis. As expected, downregulated transcriptomes dominated the changes in both *Cgas*^-/-^ (**Figure 7A**) and *Sting*^-/-^ (**Figure 7B**) cardiac neutrophils compared to WT. We identified more downregulated genes (n=2311) in *Sting*-deficient neutrophils compared to *Cgas*^-/-^ neutrophils (n=766), whereas the numbers of upregulated genes remained similar.

**Figure 7.**
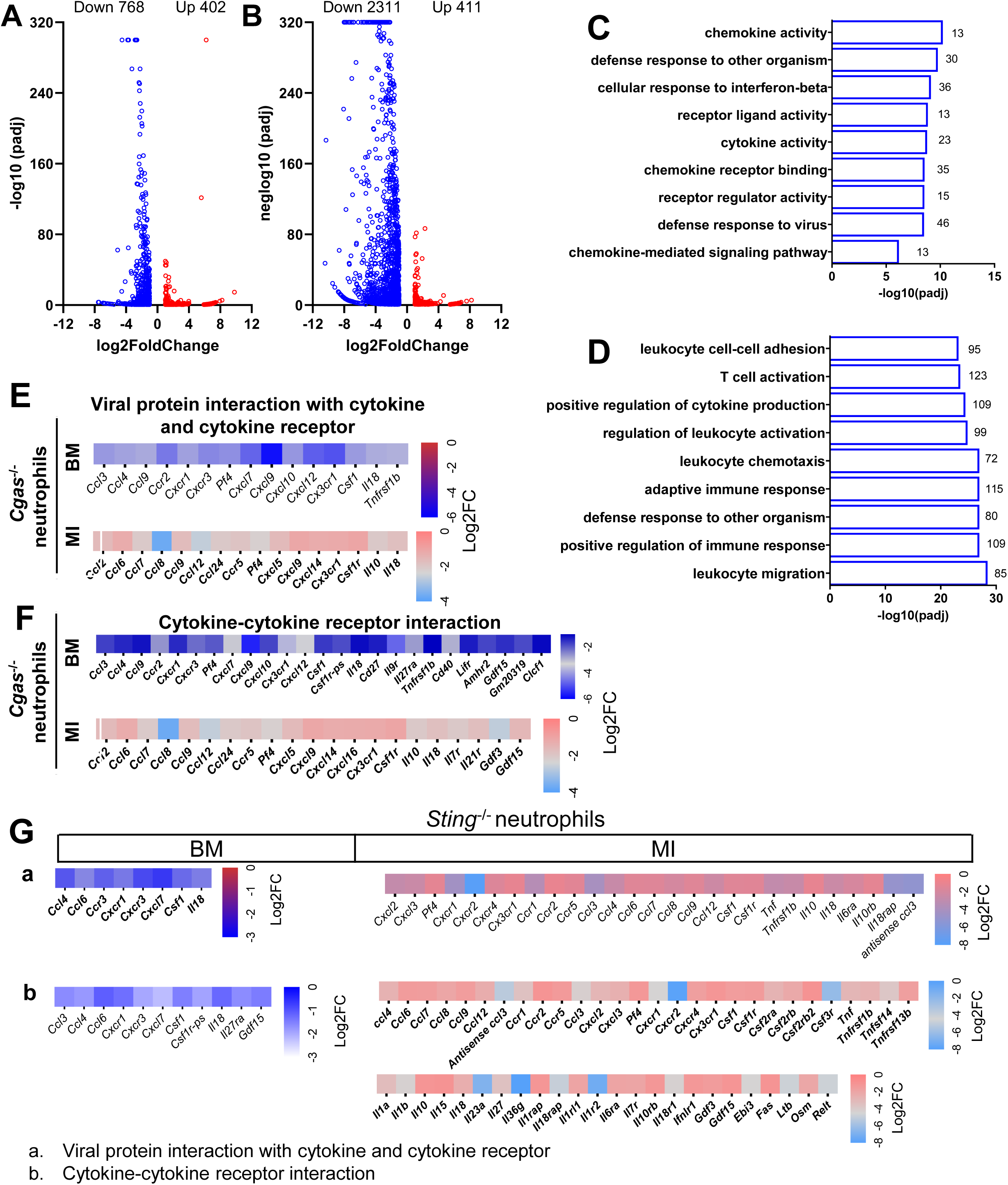
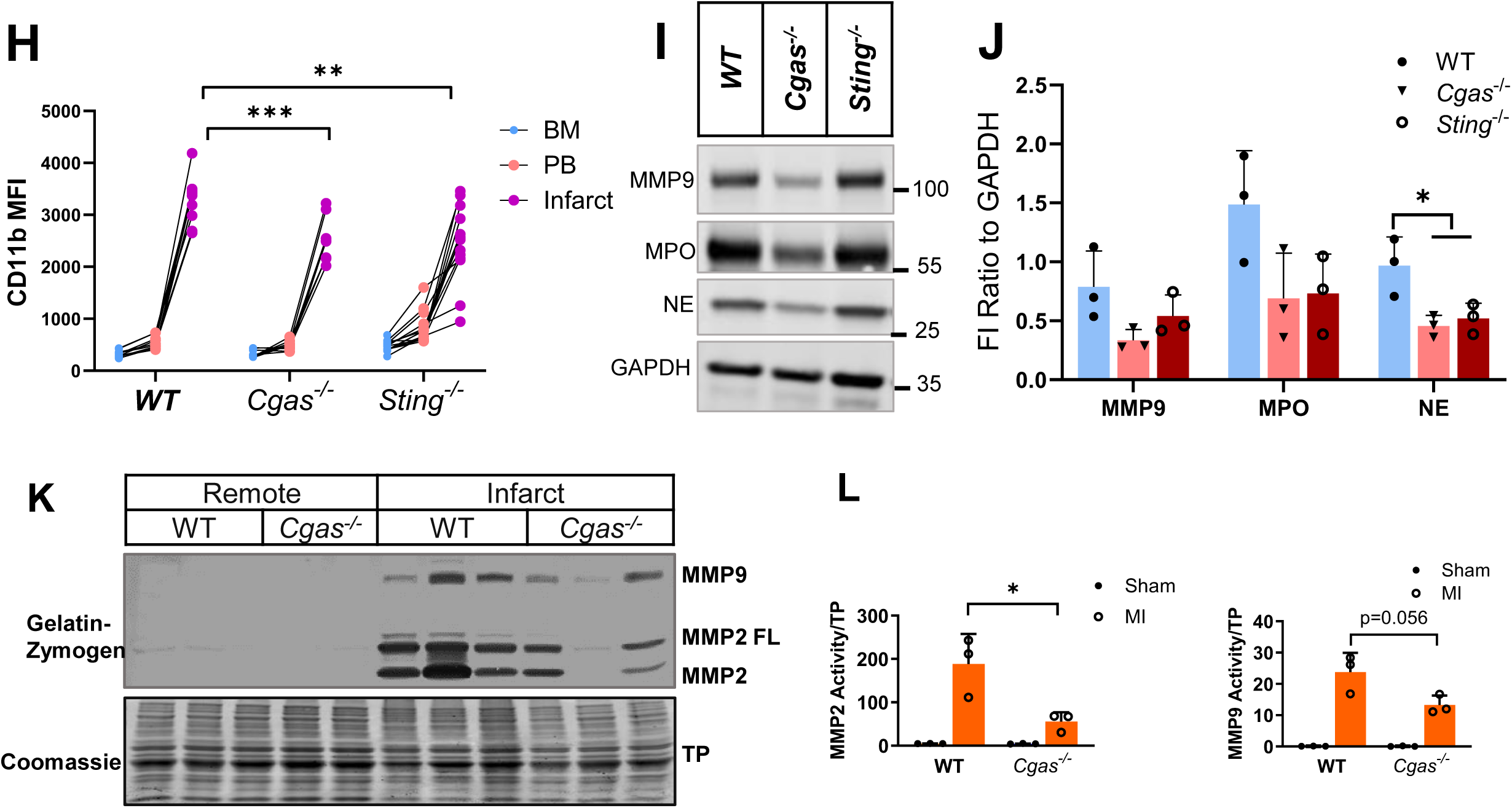
cGAS-STING pathway promotes neutrophil effector functions in the ischemic environment. Neutrophils from myocardial infarct tissue three days after MI were purified by positive selection using Ly6G beads. RNA samples purified from these neutrophils were subjected to RNA sequencing analysis. **A**-**B**. Volcano plot of differentially expressed genes in *Cgas*^-/-^ (**A**) or *Sting*^-/-^ (**B**) neutrophils compared to WT cells using criteria Log2FC greater than or equal to 2 and padj value <0.05. **C**-**D**. The top 10 most enriched GO terms in the down-regulated transcriptomes in *Cgas*^-/-^ (**C**) and *Sting*^-/-^ (**D**) neutrophils compared to WT neutrophils after MI. **E**-**G**. Heatmaps of specific genes significantly down-regulated from the indicated KEGG pathways in neutrophils from the BM and MI tissue in *Cgas*^-/-^ neutrophils compared to WT (**E**-**F**) or in *Sting*^-/-^ neutrophils compared to WT (**G**). **H**. Neutrophil activation marker CD11b was assessed from neutrophils isolated from BM, PB, and infarct of the indicated genotypes. **I**-**J**. Lysates of MI neutrophils were subjected to immunoblot analysis of neutrophil effector proteins. **K**-**L**. Lysates were prepared from remote tissue or infarct tissue after MI and subject to in-gel zymography to assess the activities of MMP2 and MMP9. *p<0.05. Supplement Figure. Modified Giemsa stain of cells from the neutrophil phase and monocyte phase after gradient centrifugation. Arrows indicate monocytes.

The top downregulated GO terms in the absence of *Cgas* (**Figure 7C**) or *Sting* (**Figure 7D**) included defense mechanisms against other organisms, chemokine and cytokine activity, response to IFNB, chemotaxis, and leukocyte activation. GO terms downregulated in cardiac neutrophils due to the loss of cGAS-STING overlapped with those in BM neutrophils (**Figures 4D** and **4F**). Intriguingly, T cell activation and adaptive immune response were GO terms uniquely downregulated in *Sting*^-/-^ neutrophils (**Figure 7D**), suggesting an unrecognized role for STING in these processes.

"Cytokine-cytokine receptor interaction" and "Viral protein-cytokine receptor interaction" were among the most enriched KEGG pathways, showing significant upregulation in WT BM neutrophils after MI compared to the sham procedure (**Figure 3F**). In neutrophils from infarct tissue, these same pathways were among the top three most enriched KEGG pathways from downregulated transcriptomes of both *Cgas*^-/-^ and *Sting*^-/-^ MI neutrophils compared to WT (**Figures 7E, 7F, 7G**). These data indicate that the decreased neutrophil functionality in the BM by cGAS-STING inhibition was maintained in the infarct environment. Interestingly, *Sting*^-/-^ MI neutrophils had more downregulated genes compared to *Cgas*^-/-^ neutrophils, suggesting STING-specific regulation of processes activated by the ischemic environment. Very few upregulated pathways and processes existed in *Cgas*^-/-^ and *Sting*^-/-^ MI neutrophils compared to WT. KEGG enrichment analysis revealed no significantly upregulated pathways for *Cgas*^-/-^ neutrophils and one for *Sting*^-/-^ neutrophils, the "Neuroactive ligand-receptor interaction."

In summary, the similar enriched pathways in cardiac neutrophils compared to BM neutrophils suggest that the cGAS-STING mediated changes during neutropoiesis in the BM persisted in the ischemic environment, connecting neutrophil priming in the BM with their function in the injury site. Our data identify predominant processes governed by cGAS-STING in neutrophils, including cytokine signaling and IFN-activated genes. The neutrophils recruited to the infarct tissue remain transcriptionally active and likely play roles in their effector function.

The transcriptomic analysis indicates that BMs from *Cgas*- or *Sting*-deficient mice generate neutrophils with dampened activation. To further determine whether this corresponds to functional changes in *Cgas^-/-^* or *Sting*^-/-^ neutrophils during acute ischemia, we first assessed neutrophil activation by measuring CD11b, a member of the β2 integrin family stored in tertiary granules and released to the cell surface upon neutrophil activation. CD11b levels remarkably increased after neutrophils migrated from BM and PB to the infarct tissue (**Figure 7H**). Loss of cGAS-STING significantly reduced CD11b on the neutrophil cell surface (**Figure 7H**), suggesting reduced degranulation and activation of these cells in response to the ischemic environment. Additionally, *Cgas-* or *Sting*-deficient neutrophils from MI tissue exhibited lower levels of matrix metalloprotease (MMP) 9, myeloperoxidase (MPO), and neutrophil elastase (NE) (**Figures 7I, 7J**). Enzyme activity of MMP9 and MMP2 in infarct tissue from *Cgas^-/-^* mice was also decreased, as demonstrated by in-gel zymography (**Figures 7K, 7L**). In summary, these data and the transcriptomic analysis support that *Cgas*- and *Sting*-deficient neutrophils have reduced functionality and a dampened response to stimulation from the ischemic environment.

## Discussion

We investigated the role of ischemia in priming neutrophils during neutropoiesis in the BM and if cGAS-STING regulates this process and neutrophil production after MI. Combining RNA sequencing of BM and MI neutrophils, cell surface marker profiling, and functional studies, we uncovered that ischemia induced a wide range of activities in BM neutrophils that are essential to their functions. The inhibition of cGAS-STING down-regulated several pathways activated by ischemia and decreased neutrophil production. Moreover, cGAS-STING governed ISG expression in neutrophils and the generation of IFIT1 neutrophils. The overlap of pathways activated by ischemia and inhibited by *Cgas* and *Sting* deletion in BM and MI neutrophils highlight neutropoiesis as an integral part of neutrophils gaining functionality and controlling neutrophil phenotype in the MI tissue. Furthermore, we identified that acute ischemia stimulates pathways essential to canonical neutrophil effector functions as well as processes involving antigen presentation, adaptive immunity, and maintenance of cell cycle-related programs. By examining neutrophils from both the origin and the destination compartments, the BM and MI tissues, respectively, the data presented provide insights into neutrophil differentiation and maturation, the connection between BM neutrophil priming and functional phenotype, and a broader functional repertoire in the setting of myocardial ischemia than previously known.

The initial response to acute myocardial ischemia involves a rapid influx of neutrophils. These cells play a crucial role in removing infarct tissue and repairing the myocardium, but their powerful weaponry also causes untoward tissue destruction. The functionality of BM neutrophils stimulated by ischemia is not well understood. However, this important aspect can determine the neutrophil phenotype in the ischemic injury site. Unlike macrophages, which rely on environmental signals for proper activation, mature neutrophils released from the BM are pre-equipped with tools formed during their differentiation and maturation. In other words, neutrophils are "made to order." This is evident from the established sequence of granules formed during neutrophil differentiation: azurophilic granules form immediately after neutrophil lineage commitment, followed by specific, gelatinase, and secretory granules^41^. These unique features of neutrophils underscore the importance of studying neutropoiesis to understand how ischemia-induced specific functions are regulated. The data presented here show that ischemia simultaneously enhances neutrophil generation and significantly upregulates phagocytosis, chemotaxis, degranulation, and cytokine-receptor signaling in BM neutrophils. Recent evidence suggests that the functional profile of neutrophils changes in response to different pathogens^42^, so defining ischemia-specific neutrophil priming in the BM will be interesting and clinically beneficial.

We identified cytokine signaling as a top KEGG pathway upregulated by ischemia in the BM. Neutrophils were reported to express few cytokine receptors limited to CXCL1 and CXCL2 at baseline^43^. However, they upregulate CCRs (CCR1 to 5) and other CXCR receptors with inflammatory stimulation^44, 45^. CCR2 and CCR5, in turn, enabled neutrophils to generate ROS and activated phagocytosis in inflammatory tissue^44, 46^. Additionally, increased CCR7 guided neutrophils to migrate to a specific destination from the inflammatory site^47^. The CXC receptors and their ligands (CXCR1, CXCR3, CXCL4, CXCL7, CXCL12, CXCL14) promoted inflammatory signaling through NFKB activation, ROS generation, enhanced phagocytosis capacity, and degranulation^43, 48^. Thus, our study suggests enhancing cytokine signaling constitutes a central pathway to prepare neutrophils to respond to ischemia.

Intriguingly, ischemia promoted pathways related to adaptive immunity, including antigen presentation and T cell function (e.g. the up-regulation of key components of the primary immunodeficiency (PID) pathway). Antigen presentation by neutrophils has only recently gathered attention. In contrast, ample evidence supports neutrophils as immune-modulating cells that suppress T-cell function in cancer. In addition, prior work had suggested neutrophils protected themselves from apoptosis by T cell receptor signaling that required Rag1^49^, a vital member of the PID pathway. Furthermore, Ischemia simultaneously enhances both survival and apoptosis programs in BM neutrophils, suggesting the life span of neutrophils, a tightly controlled process to limit the damage they can inflict, is regulated by the ischemic insult. Investigating how neutrophil life span is controlled at the level of neutropoiesis will be an interesting area for future research.

To study the role of cGAS and STING in emergent neutropoiesis after MI, we used whole-body deletion of *Cgas* and *Sting*. This was necessary since emergent neutropoiesis requires increased HSPC proliferation and commitment to the neutrophil lineage. Evidence suggests that HSPCs bypass intermediate steps to directly differentiate into lineage specific progenitors when urgent demand rises^50^. Furthermore, multiple cues coordinate emergent neutropoiesis, including transcription factors, the sensing of DAMP(PAMP) by pattern recognition receptors (PRR) on HSPCs, the stem cell niche that includes multiple types of stromal cells, and cytokines such as IL1, IL6, IFNs, and TNFa^6, 7, 50^. Using a neutrophil-specific promoter such as *Mrp8* will only delete *Cgas* and *Sting* in granulocyte/macrophage progenitors after they are committed to neutrophil lineage. This will exclude many key steps, including but not limited to HSC (hematopoietic stem cell) proliferation. In addition, HSPCs are extremely sensitive to extrinsic factors such as cytokines and alarmins and express PRRs TLR2 and TLR4. The DNA-cGAS-STING axis could be a DAMP-PRR system that promotes proliferation in HSC. A neutrophil-specific knockout strategy will likely yield limited knowledge regarding the role of cGAS-TING in neutropoiesis. Additionally, the current study is the first step in delineating cGAS-STING in neutropoiesis. Future studies using neutrophil-specific knockout animal models can help identify stages of differentiation (e.g. stem cells vs lineage-committed progenitors) influenced by cGAS-STING.

Using these models, we identified significant transcriptome changes in neutrophils deficient in *Cgas* and *Sting* from both the bone marrow (BM) and myocardial infarction (MI) tissue compared to their wild-type (WT) counterparts. The most downregulated pathway in *Cgas-* and *Sting-*deficient neutrophils was the "Cytokine and cytokine receptor signaling" pathway, which is among the most upregulated pathways in response to ischemia. Additionally, the "Nod-like receptor pathway" was prominently suppressed by *Cgas* and *Sting* deletion. The Nod-like receptor family has been implicated in neutropoiesis following MI^35, 36^. Many components of the Nod-like receptor pathway are interferon-stimulated genes (ISGs) controlled by cGAS-STING, suggesting a potential connection between this pathway and the promotion of neutrophil production after MI. Given that DAMP and PAMP sensing is a key component of emergent neutropoiesis, it is plausible that cGAS-STING serves as a PRR to detect cell-free DNA, which is elevated after MI^51^, as a signal of continuous tissue destruction to promote neutropoiesis. The decreased neutrophil production in *Cgas*^-/-^ and *Sting*^-/-^ supports this notion.

An unexpected finding of the current study is the persistent presence of PH3+ neutrophils in the infarct tissue. PH3 indicates G2 and mitosis phases in the cell cycle. Neutrophils in the infarct tissue are considered terminally differentiated and have lost the capacity to proliferate. Under inflammatory stress, progenitors committed to neutrophil lineage rapidly divide to meet the high demand and the post-mitotic maturation phase is dramatically reduced^11^. The PH3 in infarct neutrophils could be from the remnants of cell cycle machinery after rapid proliferation, and the shortened maturation time is not enough for the machinery to be degraded. Indeed, a recent proteomic analysis demonstrated cell cycle proteins were among the most upregulated proteins after the challenge of an infectious agent, and these cell cycle proteins were maintained after neutrophils migrated to the peripheral tissue^37^. The implication of this phenomenon is unclear. One study reported cell cycle proteins participated NETosis^52^. Thus, it is also possible that the cell cycle-related proteins carry out non-canonical functions. Cell cycle-related programs in the BM were gradually down-regulated while the neutrophils matured. The enhanced cell cycle programs in *Cgas*^-/-^ and *Sting*^-/-^ neutrophils could be due to the reduced cell maturity, which is supported by the data presented here.

We also found marked differences in interferon-stimulated genes (ISGs) in wild-type (WT) neutrophils compared to *Cgas*^-/-^ and *Sting*^-/-^ neutrophils after myocardial infarction (MI). Neutrophils have been shown to be highly responsive to type I interferon (IFN), with the injection of type I IFN leading to the increased expression of 325 genes in neutrophils compared to only 44 in macrophages^53^. Neutrophils with the ISG transcriptomic signature have been identified in peripheral blood (PB), spleen, and infarct tissue^11, 17, 19^. IRF3 played a key role in ISG expression in myeloid cells^19^. Expanding on this prior observation, the data presented here support the role of cGAS-STING in ISG expression. The deletion of *Cgas* and *Sting* affected more ISGs compared to *Irf3*-deficient neutrophils^19^. A possible explanation is that IRF3 is one of the two downstream targets of cGAS-STING, with the other being NFKB. In fact, most ISGs assessed in this study have transcription factor binding sites for NFKB.

While some studies using the single-cell transcriptomic approach identify the ISG neutrophils as a discreet neutrophil population^11, 17, 19^, other study suggests different ISGs were differentially expressed at specific maturation stages during neutrophil differentiation^53^. It is also not clear how the neutrophils with ISG transcriptomic signatures correlate with neutrophils with ISG protein expression. Using IFIT1 as an index marker of ISG neutrophils, we performed intracellular staining of IFIT1 to answer this question. Our data did not separate the neutrophils into IFIT1^-^ and IFIT1^+^ populations to reflect the distinct ISG^-^ and ISG^+^ groups observed in single-cell transcriptome analysis. Instead, neutrophils with a low to high IFIT1 were demonstrated by flow cytometry and IHC. One contributing factor is that *Ifit1* was not included when the ISG score was calculated, and the timing (MI day 4) also differed from the current study (MI day 3). Therefore, IFIT1+ neutrophils may have a different dynamic than the ISG-score-guided assessment^19^. Another consideration is that neutrophils with very high ISG scores may be difficult to capture because they accounted for a small percentage of the ISG^+^ neutrophils^19^. These neutrophils with ISG signature could be generated in a unique niche and captured by single-cell transcriptomic analysis, whereas IFIT1 staining depicts the overall distribution. The data presented here agree that ISG neutrophils are present in the PB before they reach the ischemic site^19^, and the higher IFIT1 in PB neutrophils is consistent with the notion that ISG expression correlates with neutrophil maturity. The function of ISG-related programs in neutrophils remains to be determined. However, *Ifnar*^-/-^ (IFNA receptor) mice had a protective phenotype in MI^24^. With the finding that neutrophils are the most responsive cells to type I IFN, it’s plausible that type I IFN-ISG is detrimental to the repair process of ischemic myocardium. Differentiating the role of neutrophil-specific IFN-ISG signaling will require neutrophil-specific knockout of the IFNA receptor in combination with conditional knockout in monocytes. However, these studies carry more weight in providing mechanistic understanding than in clinical practicality, given available cGAS-STING inhibitors are not cell type specific.

Comparisons of *Cgas*- and *Sting* deficient neutrophils indicate these two genes and their products regulate many overlapping pathways, but that STING inhibition triggered broader transcriptomic changes in the ischemic environment, perhaps indicating that MI locally activates STING-specific processes that may be type I IFN-independent^54^. Additional studies are necessary to differentiate the cGAS- and STING-specific regulation of neutrophil function.

In summary, we investigated acute myocardial ischemia-induced emergent neutropoiesis and the role of cGAS-STING in this process. We found that ischemia triggered extensive transcriptomic changes related to innate and adaptive immunity during neutrophil differentiation in the bone marrow. The enhanced functionality of BM neutrophils contributes to the phenotype of neutrophils in the ischemic myocardium. Deficiency in *Cgas* and *Sting* mitigated neutrophil production, maturation, and functionality after myocardial infarction (MI). Our data suggest that cGAS-STING may be a DAMP-sensing pathway that promotes neutropoiesis in MI. Notably, the cGAS-STING pathway governs the interferon-stimulated gene (ISG) transcriptome and the IFIT1+ neutrophil population. This study extends the previous understanding of cGAS-STING signaling in macrophages after MI. These findings are clinically relevant and may aid in translating cGAS-STING pharmacotherapies for treating acute myocardial ischemia because of the systemic nature of available pathway inhibitors. Often, research focuses on neutrophils in the infarct area. However, the evidence presented here suggests a complex process that starts in the BM. Therefore, more comprehensive studies are necessary to understand neutrophil biology and its clinical translation better.

## Materials and Methods

### Sex as a biological variable

Both male and female mice were tested initially, and we found no phenotypical differences in our model. We therefore chose one sex (male) for this study.

### Animals

Animals were kept in a pathogen-free environment with free access to food and water. They were maintained on a 12-hour light/dark cycle from 6 am to 6 pm. All procedures were approved by the Institutional Animal Care and Use Committee at the University of Texas Southwestern Medical Center. The generation of *cGas*^-/-20, 21^ and *Sting*^-/-^ mice^55^ has been described. Both knockout mice were born at normal Mendelian ratios, and they did not display overt developmental abnormalities. Further, at baseline, *cGas*^-/-^ and *Sting*^-/-^ mice manifested no cardiovascular abnormalities, with normal contractile function and blood pressure comparable to that of WT mice. All mice are from the C57BL/6J background.

### Antibodies for immunoblot and immunohistochemistry

Antibodies from Cell Signaling (Danvers, MA): rabbit anti-human cGAS (31659S), rabbit anti-STING (Cat. 13647S), rabbit anti-IRF3 (Cat. 4302), and anti-[hospho-IRF3 (Cat. 4947S). The rat anti-neutrophil antibody. Mouse anti-GAPDH was purchased from Biosynth (Gardner, MA. Cat. 10R-G109a). Anti-PH3 was obtained from Millipore Sigma (Burlington, MA. Cat. 06-570). The rat anti-CD68 (MCA19571) was from rom BioRad (Hercules, California).

### Mouse myocardial infarction model

A murine model of myocardial infarction was employed involving permanent ligation of the left anterior descending coronary artery (LAD) as described previously^23, 56^. Briefly, 10-16 weeks old *cGas*^-/-^ and *Sting*^-/-^ mice, their WT littermates, and WT animals of the same genetic background were anesthetized with 2.4% isoflurane and positioned supine on a heating pad (37°C). Tracheal intubation was performed with a 19G stump needle, and animals ventilated with room air using a MiniVent mouse ventilator (Hugo Sachs Elektronik; stroke volume, 250 μL; respiratory rate, 210 breaths per minute). Following left thoracotomy between the fourth and fifth ribs, the LAD was visualized under a microscope and ligated with 6-0 Prolene suture. The suture was placed at the LAD segment corresponding to 1.5 mm to 2 mm below the lower edge of the left atrium. Regional ischemia was confirmed by visual inspection of discoloration of myocardium distal to the occlusion under a dissecting microscope (Leica). Sham-operated animals underwent the same procedure without occlusion of the LAD. After LAD ligation, hearts were collected at 24-, 48-, 72-hour, and 1-week. Left ventricles were separated into remote and infarct-at-risk zones for downstream analysis.

### Bone marrow neutrophil isolation

Mice were euthanized and the surface is sprayed with the 70% ethanol. The soft tissues were removed from the femurs with a sterile scalpel and the clean bones (*n* = 4) were transferred into a petri dish on ice. Both ends of the long bone (epiphysis) of the femur were cut to expose the bone marrow (BM). The PBS was used to flush out the bone marrow with and collected in a 15 mL tube. The BM cell suspension was centrifuged at 300 g for 5 min, and the pellet was resuspended in cell sorting buffer containing 2% FBS and 1mM EDTA and filtered using a 70µm cell strainer immediately before isolation. We use three methods to purify neutrophils. 1) Fluorescence-activated cell sorting (FACS). We label the BM single cell suspension with CD45, CD11b, Ly6G, CCR2, and Ly6C. The CD45^+^CD11b^+^Ly6G^+^Ly6C^int^ cell population (neutrophils) is sorted and collected for down stream analysis. 2) Gradient centrifugation. The neutrophils are isolated per published protocol^57^. The resulting bone marrow cells are collected and washed with RPMI 1640 1X supplemented with 10% FBS and 2 mM EDTA for 7 minutes at 427 × g at 4°C. The neutrophils were purified by density gradient method^57^ with some modification. The density gradient was prepared by adding 3 ml of Histopaque 1119 (density, 1.119 g/ml) in a 15-ml conical centrifuge tube, followed by overlaying with 3 ml of Histopaque 1077 (density, 1.077 g/ml). The bone marrow cell suspension is slowly transferred on top of the Histopaque 1077 layer. The grandient-cell column is centrifuged for 30 minutes at 872 × g at room temperature without brake. After centrifugation, the neutrophils is located at the interface of the two gradient layers. The neutrophils are washed with RPMI 1640 1X supplemented with 10% FBS and 1% penicillin/streptomycin by centrifuging at 1400 rpm for 7 minutes at 4°C. The neutrophils are processed according protocols of downstream analysis. 3). Ly6G Immunomagnetic separation (IMS). The BM cell suspension prepared as outlined above was incubated with anti-mouse Ly6G-Biotin antibody (Bioledgend, Cat. 27604) for 10 minutes at room temperature. After washing with sorting buffer, the cells were resuspended and incubated with Magnisort^TM^ Steptavidin Beads (Invitrogen, Cat. MSPB-6003-74) 10 minutes. The Ly6G labeled neutrophils in 2.5 ml sorting buffer were transferred to a 5ml tube followed by inserting the tube into the magnet for 5 minutes at room temperature. The supernatant is removed and the resulting cell pellet is washed for additional two more times with the sorting buffer. The Ly6G positive cells were collected and stored for down stream assays. When indicated, the non-neutrophil layer, which is enriched with monocytes, was collected and used as positive controls in immunoblot analysis of the cGAS-STING expression.

### Cardiac neutrophil isolation

Cardiac immune cell isolation as performed based on our published protocol^23, 56^. Briefly, The MI tissue were separated from the remote myocardium and rinsed in 4°C PBS. The tissue was extensively rinsed to remove circulating blood followed by mincing with surgery scissors. The minced tissues were then incubated at 37°C for 45 mins in a Thermomixer (Eppendorf) with shaking set at 120rpm in the digestion buffer that contained the following enzymes (all from Sigma, St. Louis, MI): collagenase I (cat. no. SCR103,) 450 unit/mL, hyaluronidase (cat. No. H3884) 60 unit/ml, DNase (cat. 9003-98-9) 60 unit/ml. The tissues were then triturated and passed through a 40 μm strainer. Cells were then washed in 50ml 4°C PBS containing 1% Fetal Calf Serum (FCS) by centrifugation 10 minutes at 400 g. The resulting cell pellet was resuspended in cell sorting buffer before FACS, flow cytometry, or immunomagnetic isolation using the isolation kit from Miltenyi (cat.130-120-337). For FACS, the neutrophils population is defined as CD45^+^CD11b^+^CCR2^-^ Ly6C^int^Ly6G^+^.

### Flow cytometry

The single cell suspension from the infarct tissue, peripheral blood (PB), and bone marrow (BM) was obtained and the cell concentration was adjusted to 1×10^6^/100 μL with sorting buffer. The cells were incubated first with purified anti-mouse CD16/32 (FcBlock) for 10 min at 4 °C followed by adding antibodies to the concentration specified in the table. The cells were incubated with antibodies at 4 °C for 15 min. After the incubation, the cells were pelleted by centrifugation at 500g for 5 minutes and washed two times in the sorting buffer. The labeled cells were subjected to scanning by flow cytometer. Gating strategies are described in the Results section. Mean fluorescence intensity was calculated as the median of the indicated fluorescent parameter using FlowJo software. Cell sorting was performed on an Aria III Fusion (BD Biosciences). No sodium azide in the FACS buffer for sorting experiments. Antibodies used for flow cytometry: from Biolegend: CD45-PE/Cy7 (cat# 103114), CD11b-PB (cat# 101223), F4/80-PE (cat#123109), Ly-6G-APC (cat# 127613), Ly-6C-BV711 (cat# 128037), CD206-PerCP/Cy5.5 (cat# 141715), CD64-FITC (cat# 164405), CCR2-BV421 (cat# 150605), CD182 APC-cy7 (cat# 149313), CD62L-Kiravia blue-520 (cat# 104463), CD184-BV605 (cat# 153805). From ThermoFisher: Siglec F-PE (cat#12B-1702-82), CD101-PE (cat# 12-1011-82).

### RNA isolation from neutrophils

We used Quick-RNA miniprep Kit from Zymo Research (Irvine, CA) for RNA isolation according to the instruction from the manufacturer. The RNA isolated using this method contains RNAs that equals or greater 17 nt, which include small/microRNAs. The bone marrow neutrophils used for RNA isolation wwere purified by FACS. A total of 5×10^5^ neutrophils from each mouse were used. The cardiac neutrophils were isolated using the Neutrophil Isolation Kit (Miltenyi Biotec, Gaithersburg, MD. Cat. 130-097-658). The infarct tissues of three MI hearts collected three days after LAD ligation were pooled together for each data point. Neutrophils (∼3×10^5^) from the pool MI tissue were used for RNA isolation. We added RNase inhibitor (SUPERase•In™ RNase Inhibitor, Cat. AM2696, Thermofisher, Waltham, MA) at a concentration of 0.2 u/ul in RNA Prep Buffer and RNA Washing Buffer. The concentration and quality of the isolated RNA are measured by NanoDrop and freezed in -80 for downstream analysis.

### Quantitative RT-PCR

We use 250 to 500 ng RNA for the reverse transcription reaction using SuperScript (ThermoFisher, Waltham, MA). The resulted cDNA was diluted 5x, and 2 μL of the final cDNA solution is used in the subsequent quantitative polymerase chain reaction analysis (qPCR) with Sybergreen according to manufacturer’s instructions (Roche, Indianapolis, IN). Primer sequences are as follows.

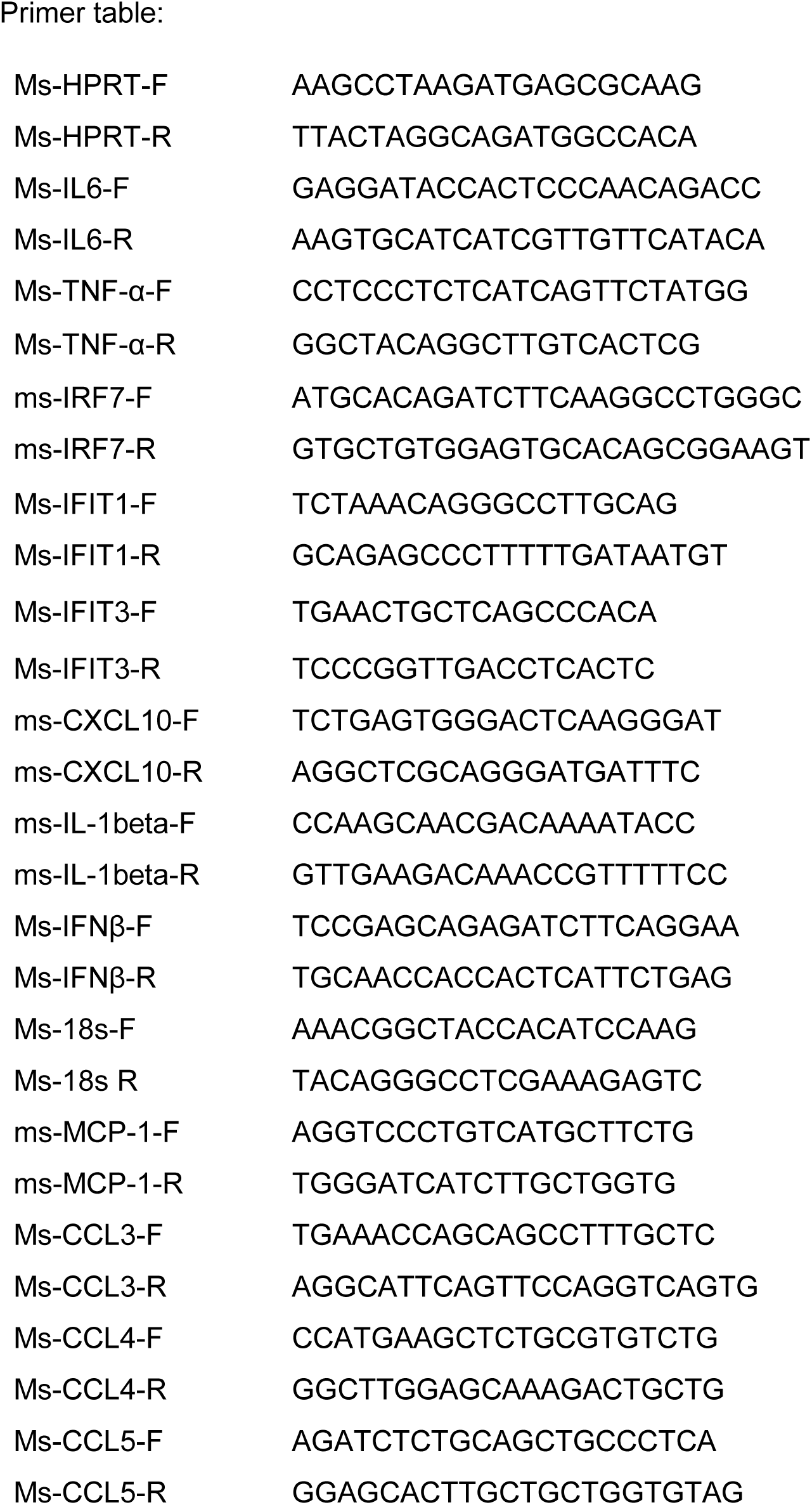
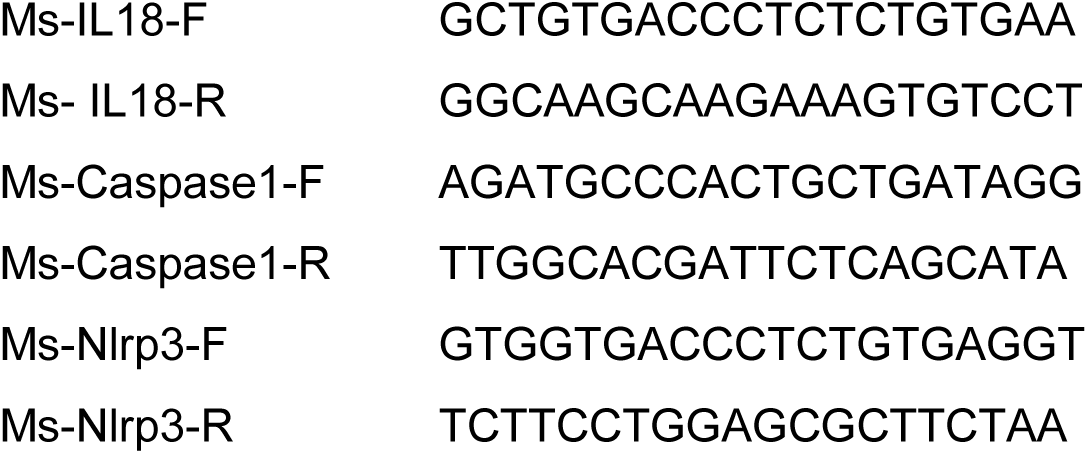

### Neutrophils RNA sequencing

RNA sequencing was performed by Novogene America (Sacramento, CA). Raw reads of fastq format were firstly processed through in-house perl scripts. In this step, clean data (clean reads) were obtained by removing reads containing adapter, reads containing ploy-N, and low-quality reads from raw data. All the downstream analyses were based on the clean data with high quality. Reference genome (mm10) and gene model annotation files were downloaded from genome website directly. Index of the reference genome was built using Hisat2 v2.0.5 and paired-end clean reads were aligned to the reference genome using Hisat2 v2.0.5. Then, the abundance of each transcript was quantified using feature Counts v1.5.0-p3. Differentially expressed genes (DEGs) analysis was performed using the DEseq2 package, and gene expression was normalized using the relative-log-expression (RLE) in DESeq2.30 Genes with an adjusted P-value <.05 found by DESeq2 were assigned as differentially expressed.

For gene ontology (GO) analysis, metascape (http://metascape.org) was employed to perform gene enrichment and functional annotation analyses in neutrophils with the following ontology sources: Kyoto Encyclopedia of Genes and Genomes (KEGG) Pathway, GO Biological Processes, and Reactome Gene Sets.31 GO terms with corrected P-value <.05 were considered significantly enriched by differential expressed genes. KEGG analysis was used cluster Profiler R package to test the statistical enrichment of differential expression genes in KEGG pathways. For genomic integrity analysis (copy number variation [CNV] analysis) was referred to a previous protocol. Preparation of RNA library and transcriptome sequencing was conducted by Novogene Co., LTD (Beijing, China). Genes with adjusted p-value < 0.05 and |log2(FoldChange)| > 0 were considered as differentially expressed.

### Identify the interferon responsive neutrophil population by intracellular IFIT1 staining and flow cytometry

Cell suspensions from MI tissue, PB and BM were prepared from the same mice three days after MI. PB (300 µl) was collected via retro-orbital bleeding and diluted in 1.5 ml Hanks’ balanced salt solution (HBSS) containing 15 mM EDTA. The resulting cell suspension was centrifuged at 500g for 10 min at 4°C. Red blood cells were removed by incubating the cells with 5 ml ammonium-chloride-potassium (ACK) lysis buffer (Biolegend) for 5 min at room temperature followed by adding 10 ml RPMI supplemented with 3% FBS. The remaining cells were collected by centrifugation at 500*g* for 5 min and washed twice with 10 ml HBSS with 2 mM EDTA and 2% BSA. The cells were incubated in flow buffer containing 1% BSA and 1%FBS in PBS with blocking agent anti-mouse CD16/CD32 for 10 minutes on ice. Cell suspension from BM was washed with PBS twice followed by anti-mouse CD16/CD32 blocking. The cells prepared from PB and BM were stained with PE/cy7-conjugated anti-CD45, PB-conjugated anti-CD11b, APC-conjugated anti-Ly6G and BV711-conjugated anti-CXCR4 antibodies were added and incubated for 20 min at 4°C protected from light. After incubation, cells were washed with PBS twice followed by fixation and permeablization in 1 ml PBS containing 2% PFA (Electron Microscopy Sciences) and 0.1% saponin (Sigma–Aldrich) at 4 °C for 30 min. The cells were collected by centrifugation at 1,500g for 5 min at 4 °C and washed with 1 ml Wash Buffer (PBS containing 0.2% BSA and 0.1% saponin) before blocking with 5% goat serum and 5% BSA for 30 min. The cells were stained with anti-IFIT1/p56 antibody (Sigma– Aldrich) for 30 min at 4 °C in 200 µl staining buffer (PBS containing 5% goat serum, 5% BSA and 0.1% saponin). After washing the cells twice with 1 ml Wash Buffer, the cells were incubated with secondary goat anti-rabbit-Alexa Fluor 488 antibody (Invitrogen) for 30 min before washing and resuspending the cells in the flow buffer. The stained cells were analyzed on a Flow Cytometer (BD). Fixation, washing, staining and sorting were performed at a concentration of 5–10 × 106 cells per ml.

### Neutrophil Respiratory Burst Assay

We performed the assay using the Neutrophil/Monocyte Respiratory Burst Assay Kit (#601130, Caymen) according to manufacturer’s instruction. Briefly, peripheral blood (PB) is obtained using the submandibular blood collection method. After puncturing the vein with a 25 gauge needle, 100 µl of PB is collected into 1.5 ml Eppendorf tube that is rinsed with heparin prior to the blood collection. The the PB samples are incubated with 0.5 µg/ml dihydrorhodamine 123 (DHR123) for 15 min at 37°C followed by adding PMA (200 µM final concentration) and additional 45 minutes of incubation. After the incubation is completed, the red blood cells (RBC) is removed by RBC lysis buffer and the leukocyte pellet is collected by centrifugation at 500g for 10 minutes. The leukocytes are labeled with CD45 and Ly6G on ice for 10 minutes before subjecting to flow cytometry analysis.

### Immunoblot analysis

The protocol for immunoblot analysis was as described previously24 with minor modifications. Briefly, tissues from the remote and infarct/risk areas of myocardial tissue were homogenized using a Dounce homogenizer containing T-PER lysis buffer from ThermoFisher (Waltham, MA) supplemented with protease inhibitors and phospho-STOP (Roche, Indianapolis, IN). Tissue lysates were centrifuged at 12,000g for 12 minutes. The supernatant was collected, and the protein concentration was measured by Bradford assay (Bio-Rad, Hercules, CA). 15 μg of total protein was loaded onto an SDS-PAGE gel and ultimately transferred to a nitrocellulose membrane. The membrane was blocked with 5% BSA, followed by incubation with primary antibodies (overnight, 4°C) and secondary antibodies (1h, RT). The membrane was scanned and quantified using an Odyssey scanner (LI-COR, Lincoln, NE).

### cGAMP stimulation

Neutrophils isolated by density gradient centrifugation are seeded on 12-well cell culture dishes by Corning (Corning, NY) at 5×10^5^ per well in RPMI media supplemented with 10% FBS and Penicillin-Streptomycin. The cells are cultured at 37°C and 5% CO2 for one hour before adding cGAMP or the control compound of cGAMP at the concentration of 15 μM for three hours. After the stimulation, the cells are collected by centrifugation and washed with PBS at 400g for 5 minutes at 4°C. The cells collected are used to extract total protein for immunoblot analysis and for RNA isolation for RT-qPCR. using the Quick-RNA miniprep Kit from Zymo Research.

### Immunohistochemistry

Hearts harvested at different time points after LAD ligation were fixed in 4% PFA for 48 hours. Transverse sections at the following levels were prepared: above the suture, at the suture, and 0.5 mm, 1 mm, 1.5 mm, 2 mm, 2.5 mm distal to the suture. Detection of neutrophil, macrophages, myofibroblasts, and microvessels was performed according to the following protocol. Sham- or MI-operated mice at indicated time points were anesthetized with isoflurane. Immediately after cervical dislocation, the heart was exposed and the right atrium was nicked with a scalpel, and 10 mL ice-cold PBS was injected into the left ventricular cavity. The heart was then perfused with ice-cold 4% paraformaldehyde in PBS. After trimming away the atria and great vessels, the heart was embedded in OCT compound (VWR International) and frozen in isopentane chilled in liquid nitrogen. The frozen tissue sections were obtained in transverse orientation starting at the level of LAD suture and progressing toward the apex at 0.5 mm intervals. Sections were incubated in PBS containing 0.1% Triton-X (5 min, RT), and non-specific antibody binding sites were pre-blocked with PBS plus 5% BSA. Sections were then incubated with primary antibodies in PBS with 5% BSA (overnight, 4°C) in a humidified chamber. After rinsing with PBS 5 times, the sections were next incubated with appropriate fluorophore-conjugated secondary antibodies (1h, RT). Stained sections were mounted with anti-fade mounting medium before proceeding to fluorescence microscopy.

### Echocardiography

Mouse echocardiograms were recorded on conscious, gently restrained mice using a Vevo 2100 system and an 18-MHz linear probe as described.^23, 56^. A short-axis view of the LV at the level of the papillary muscles was obtained, and M-mode recordings were obtained from this view. Left ventricular internal diameter at end-diastole (LVIDd) and end-systole (LVIDs) were measured from M-mode recordings. Fractional shortening was calculated as (LVIDd − LVIDs)/LVIDd (%).

### Statistical analysis

Findings are expressed as mean ± SD. The data were analyzed using statistical software (GraphPad Prism, version 7.01; GraphPad Software, San Diego, CA). An unpaired Student t test was performed to analyze 2 independent groups. One-way ANOVA coupled with the Tukey post-hoc test was used for pairwise comparisons. In representative datasets, we have also employed nonparametric tests (Wilcoxon rank sum test, Wilcoxon two-sample test, Kruskal-Wallis test). A value of p < 0.05 was considered statistically significant and results are depicted throughout as follows: *p<0.05; **p<0.01, ***p<0.001.

## Acknowledgement.

1. J. Z., X. L., and D.J.C are supported by National Heart, Lung, and Blood Institute award R01HL145298, VA MERIT Review award 1 I01 BX004562, and 1 P01 HL160488-01A1. M.C.M. is supported by the National Heart, Lung, and Blood Institute of the National Institutes of Health by award number T32HL125247. P. P. is supported by 1 P01 HL160488-01A1.

